# Establishing *Physalis* as a new *Solanaceae* model system enables genetic reevaluation of the inflated calyx syndrome

**DOI:** 10.1101/2022.07.29.502011

**Authors:** Jia He, Michael Alonge, Srividya Ramakrishnan, Matthias Benoit, Sebastian Soyk, Nathan T. Reem, Anat Hendelman, Joyce Van Eck, Michael C. Schatz, Zachary B. Lippman

**Author notes:** Ohalo Genetics, Aptos, CA 95003, USA (M.A.); LIPME, Université de Toulouse, INRAE, CNRS, Castanet-Tolosan 31326, France (M.B.); Center for Integrative Genomics, University of Lausanne, CH-1005 Lausanne, Switzerland (S.S.); Benson Hill, St. Louis MO 63132 (N.T.R).

## Abstract

The highly diverse *Solanaceae* family contains several widely studied model and crop species. Fully exploring, appreciating, and exploiting this diversity requires additional model systems. Particularly promising are orphan fruit crops in the genus *Physalis,* which occupy a key evolutionary position in the *Solanaceae* and capture understudied variation in traits such as inflorescence complexity, fruit ripening and metabolites, disease and insect resistance, self-compatibility, and most notable, the striking Inflated Calyx Syndrome (ICS), an evolutionary novelty found across angiosperms where sepals grow exceptionally large to encapsulate fruits in a protective husk. We recently developed transformation and genome editing in *Physalis grisea* (groundcherry). However, to systematically explore and unlock the potential of this and related *Physalis* as genetic systems, high-quality genome assemblies are needed. Here, we present chromosome-scale references for *P. grisea* and its close relative *P. pruinosa* and use these resources to study natural and engineered variation in floral traits. We first rapidly identified a natural structural variant in a *bHLH* gene that causes petal color variation. Further, and against expectations, we found that CRISPR-Cas9 targeted mutagenesis of 11 MADS-box genes, including purported essential regulators of ICS, had no effect on inflation. In a forward genetics screen, we identified *huskless*, which lacks ICS due to mutation of an *AP2-like* gene that causes sepals and petals to merge into a single whorl of mixed identity. These resources and findings elevate *Physalis* to a new *Solanaceae* model system, and establish a new paradigm for the search of factors driving ICS.

## INTRODUCTION

The *Solanaceae* is one of the most important plant families in fundamental and applied research due to its remarkable morphological and ecological diversity, and its far-reaching economic value from its many members used as food crops, ornamentals, and sources of pharmaceuticals (Añibarro-Ortega et al., 2022; Gebhardt, 2016; Shenstone et al., 2020) . The most studied *Solanaceae* include major food crops eggplant (*Solanum melongena*), pepper (*Capsicum annuum*), potato (*Solanum tuberosum*), and tomato (*Solanum lycopersicum*), in addition to model species petunia (*Petunia hybrida*) and tobacco (*Nicotiana benthamiana*). However, various species-specific limitations have made tomato the preferred model with a full suite of genetic and genomic resources that enable maximal biological discovery and translation to agriculture.

Developing new *Solanaceae* model systems that equal the utility of tomato is essential to study incompletely explored diversity, including traits of economic importance. Most challenging is identifying potential systems with noteworthy comparative and species-specific variation that, critically, can be dissected by efficient forward and reverse genetics, enabled by tractable genomics, genome editing, and cultivation. We previously identified species in the genus *Physalis* as promising in all these aspects (Lemmon et al., 2018) . The genus includes ornamental and orphan crops such as Chinese lantern (*P. alkekengi*), tomatillo (*P. philadelphica* and *P. ixocarpa*), goldenberry (*P. peruviana*), and groundcherry (*P. grisea* and *P. pruinosa*), and many other species yielding edible fruits. *Physalis* also occupies a key phylogenetic position that complements other *Solanaceae* models and captures the largest species richness in the tribe Physalideae, which itself has the highest diversity of *Solanaceae* genera (Deanna et al., 2019; Pretz & Deanna, 2020; Zamora-Tavares et al., 2016) . Moreover, *Physalis* species show substantial variation in developmental and molecular traits, including inflorescence complexity, secondary metabolism, and disease resistance (Baumann & Meier, 1993; Huang et al., 2020; Park et al., 2014; Whitson, 2012; W.-N. Zhang & Tong, 2016). However, the most conspicuous and impressive feature of *Physalis*, also found also in other angiosperms, is the Inflated Calyx Syndrome (ICS), a remarkable evolutionary novelty where sepals grow excessively large after fertilization to form balloon-like husks that encapsulate fruits (He et al., 2004; Wilf et al., 2017).

Dissecting the evolutionary and mechanistic origins of morphological novelties is a fundamental goal in biology (Muller & Wagner, 1991; Shubin et al., 2009), and it is not surprising that botanists and evolutionary biologists have long been fascinated by ICS (He et al., 2004; U. T. Waterfall, 1958; Wilf et al., 2017) . Though *Physalis* has historically lacked molecular and functional genetics tools, studies on ICS over the last few decades have suggested a central role for two MADS-box genes, including an ortholog of one gene in potato, *STMADS16* (ortholog of Arabidopsis *AGAMOUS-LIKE 24*), that causes leaf-like sepals when over-expressed in other *Solanaceae* (He et al., 2004). Prompted by this observation, supportive molecular and functional genetic data generated within *Physalis* suggested that heterotopic expression of the *STMADS16* ortholog *MPF2* was key to the evolution of ICS. Later studies suggested this essential role emerged from modified *cis-*regulatory control of *MPF2* by the *euAP1-like* gene *MPF3* (He & Saedler, 2005; Zhao et al., 2013). A recent genome of *P. floridana* and additional functional work suggested that loss of another MADS-box gene, *MBP21/JOINTLESS-2* (*J2*), was also critical, and seemingly reinforced an additional conclusion that fertilization is an integral physiological driver of ICS (Lu et al., 2021). The proposed role of fertility and previous findings that flower-specific *MPF2* expression is ancestral to ICS suggested this trait may have been lost during evolution (He & Saedler, 2007; Hu & Saedler, 2007). However, a recent deeply sampled taxonomic study of the Physalideae showed that ICS was gained multiple times in a stepwise and directional manner, from non-inflation to enlarged sepals appressed to the fruit (accrescent-appressed), and finally to an inflated calyx (Deanna et al., 2019) . These findings, along with independent emergence of ICS in other angiosperms (Deanna et al., 2019), may indicate deeper genetic and molecular complexity behind ICS, determined by factors besides *MPF2* and other proposed mediating *MADS-box* genes (Deanna et al., 2019; Hu & Saedler, 2007).

Outstanding questions on ICS and our broad interest in *Solanaceae* biology and agriculture led us several years ago to begin establishing *Physalis* as a new model system. We developed efficient *Agrobacterium*-mediated transformation and CRISPR-Cas9 genome editing in the diploid groundcherry species *P. grisea*, and demonstrated the utility of these tools by mutating orthologs of tomato domestication genes in groundcherry to improve productivity traits (Lemmon et al., 2018; Swartwood & Van Eck, 2019) . More recently, *P. grisea* was critical in revealing pleiotropic functions of an ancient homeobox gene, and in dissecting evolution of redundancy between duplicated signaling peptide genes controlling stem cell proliferation in the *Solanaceae* (Hendelman et al., 2021; Kwon et al., 2022) . However, high-quality reference genomes of *P. grisea* and other species have been lacking, and are needed to promote the full potential and deployment of this system as has been achieved in tomato. Here, we report high-quality chromosome-scale genomes for *P. grisea* and its close relative *P. pruinosa.* We demonstrate the power of these resources in enabling forward and reverse genetics, by revealing multiple genotype-to-phenotype relationships in floral development, including ICS. Our work establishes *Physalis* as a new *Solanaceae* reference system that can advance comprehensive studies of longstanding and emerging biological questions within and beyond the genus.

## RESULTS

### Chromosome-scale reference genomes of *P. grisea* and *P. pruinosa*

Among *Solanaceae* genera*, Physalis* is more closely related to *Capsicum* (pepper) than *Solanum* (e.g., eggplant, potato, tomato) (**Figure 1A**). Chinese lantern, tomatillo and many other *Physalis* orphan crops are self-incompatible large plants with tetraploid genomes, making them challenging to develop into model systems. In contrast, the groundcherry species *P. grisea, P. pruinosa*, and close relatives have reasonable genome sizes (estimated ∼1-2 Gb), are diploid, self and cross compatible, have rapid generation times (first mature fruit 66-70 days after sowing), and are easy to grow and manage in both greenhouses and fields. The taxonomy and naming of *Physalis* species has a convoluted past that was recently clarified (Pretz & Deanna, 2020) . *P. pruinosa* was initially designated to describe *Physalis* in the northeastern United States showing erect or prostrate growth, with large, thick and coarsely sinuate-dentate leaves (Rydberg, 1896). A revision of *Physalis* in the last century proposed *P. pubescens* var. *grisea* to differentiate species included in *P. pruinosa* (U. T. Waterfall, 1958). Additional species were then identified (U. T. Waterfall, 1967), and *P. pubescens* var. *grisea* was ultimately recognized as a separate species, *P. grisea* (Martínez, 1993; Pretz & Deanna, 2020).

**Figure 1.**
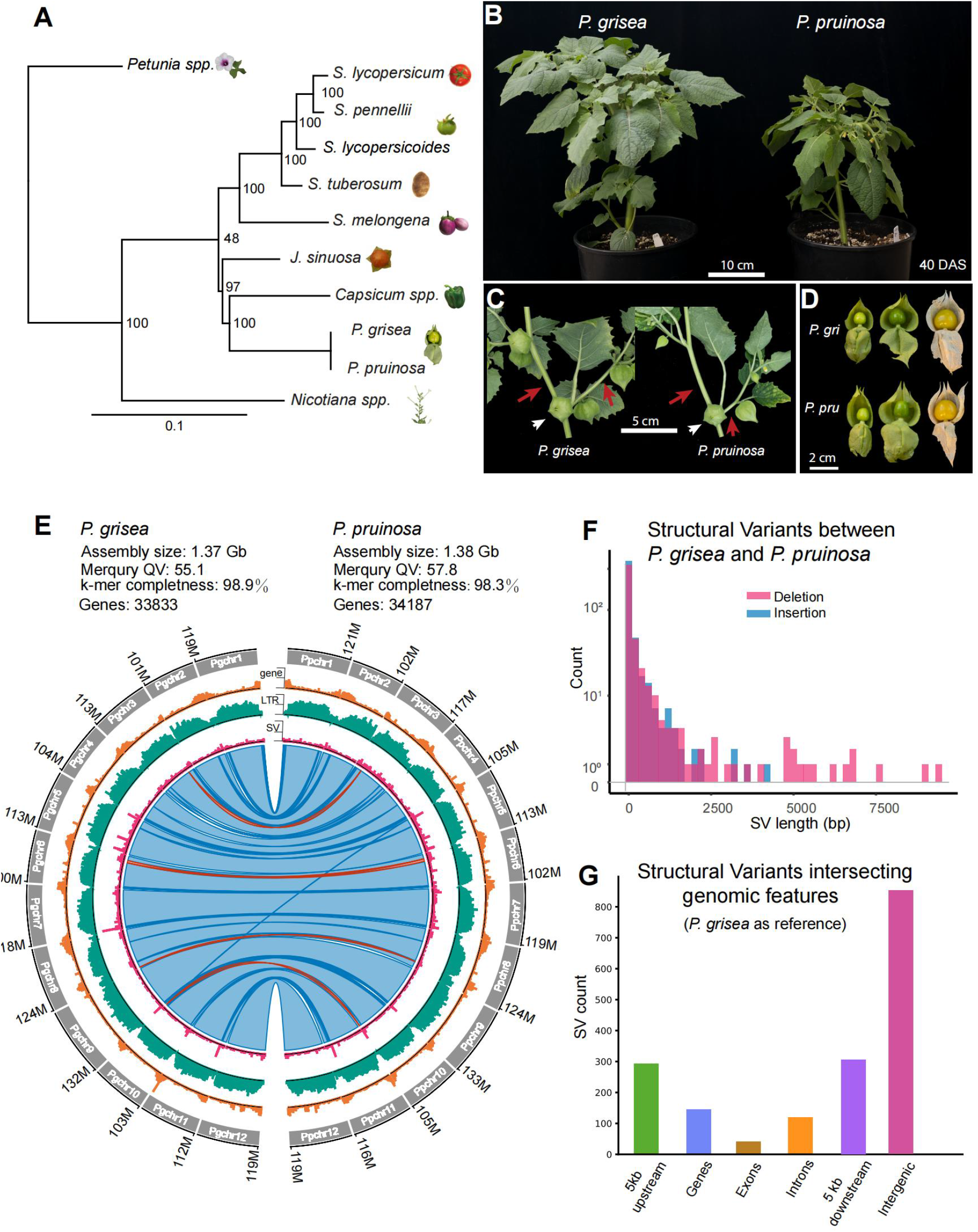
Reference-quality genome assemblies of *P. grisea* and *P. pruinosa*. **A.** Phylogeny of selected *Solanaceae* species based on the 20 most conserved protein sequences (see **Methods**). **B.** Whole plant images of *P. grisea* and *P. pruinosa* 40 days after sowing (DAS) in greenhouse conditions. **C.** Sympodial shoot architectures of *P. grisea* and *P. pruinosa*. Quantification of internode lengths is in **Source Data 2**. **D.** Images of *P. grisea* and *P. pruinosa* calyces and fruits at different stages. Husks were manually opened to show fruits **E.** Circos plots comparing *P. grisea* and *P. pruinosa* genomes. Circos quantitative tracks are summed in 100 kbp windows and show number of genes (lower tick = 0, middle tick = 25, higher tick = 49), LTR retrotransposons (lower tick = 0, middle tick = 102, higher tick = 204) and SVs (lower tick = 0, middle tick = 4, higher tick = 9). The inner ribbon track shows whole genome alignments, with blue indicating forward-strand alignments and red indicating reverse-strand alignments (inversions). Darker colors indicate alignment boundaries. **F.** Distribution of deletion and insertion SVs between 30 bp and 10 kbp from *P. pruinosa* compared to *P. grisea*, summed in 200 bp windows. **G.** Counts of SVs intersecting genomic features comparing *P. pruinosa* to *P. grisea*.

As *P. grisea* and *P. pruinosa* are closely related, they share similar vegetative and reproductive shoot and organ morphologies, including inflated calyxes encapsulating fruits of similar size, shape, and color (**Figure 1B-D**). Their primary shoots terminate in a single flower inflorescence after 5-6 leaves, and new shoots emerge according to the sympodial growth habit that is characteristic of all *Solanaceae* (Lemmon et al., 2018) . In *Physalis,* sympodial units are comprised of one leaf, one flower and two axillary (sympodial) shoots (**Figure 1C**). A conspicuous feature distinguishing *P. pruinosa* from *P. grisea* is the absence of purple pigmentation on stems and petal nectar guides. *P. pruinosa* also has narrower leaves and a smaller stature due to shorter internodes (**Figure 1B, D; Source Data 2**).

Based on the features described, *P. grisea* and *P. pruinosa* are excellent candidates to occupy a key phylogenetic position among *Solanaceae* model systems. We integrated PacBio high fidelity (HiFi) and Oxford Nanopore (ONT) long-read sequencing to establish highly accurate and complete chromosome-scale genome assemblies for both species with assembly sizes of 1.37 Gb and 1.38 Gb, respectively (**Figure 1E**). The *P. grisea* and *P. pruinosa* assemblies are the first *Physalis* genus reference-quality assemblies, demonstrating substantially improved contiguity, accuracy, and completeness compared to a recent *P. floridana* genome (Lu et al., 2021) (**Supplemental Table S1**). Specifically, the *P. floridana* genome has an error rate (errors/bp) of 3.83x10^-4^ and a contig N50 of 4.6 Mbp, whereas our assemblies produced substantially lower error rates of 3.09x10^-6^ (*P. grisea*) and 1.66x10^-6^ (*P. pruinosa*) and much higher contig N50s of 31.6 and 82.2 Mbp, respectively, with gapless assemblies of chromosomes 5 and 7 for *P. pruinosa*.

Based on RNA-sequencing data from vegetative and reproductive tissues ((Lemmon et al., 2018), and **Methods**), we annotated 33,833 and 34,187 genes in the *P. grisea* and *P. pruinosa* assemblies, respectively, with most genes concentrated at the ends of the 12 chromosomes, as is observed in other *Solanaceae* genomes (Kim et al., 2014; Sato et al., 2012; Wei et al., 2020; X. Xu et al., 2011) (**Figure 1E**, see **Methods**). Both genomes are highly repetitive, with 79% of sequence representing transposable elements, especially LTR retrotransposons (**Figure 1E**). Comparing the two genomes, we observed nearly complete macrosynteny across all 12 chromosomes, consistent with the close relationship of these species, but also detected a few small-scale inversions and translocations (**Figure 1E**). Calling single nucleotide polymorphisms (SNPs) using *P. pruinosa* Illumina short read sequences against the *P. grisea* reference revealed 60,087 homozygous SNPs, with predicted high impact changes (SNPeff, (Cingolani et al., 2012)) on 43 gene transcripts (**Supplemental Table S3**). Despite the broad similarity of these genomes, we identified over 900 structural variants (SVs) between 30 bp and 10 kbp, many of which intersect coding and putative *cis*-regulatory sequences (**Figure 1F, G and Supplemental Table S4, S5**). Some of these variants could explain phenotypic differences between *P. grisea* and *P. pruinosa*.

### A structural variant in the bHLH transcription factor gene *ANTHOCYANIN1* controls nectar guide color variation

We first sought to utilize our genomes to map the most conspicuous phenotype distinguishing the two species, nectar guide color variation*. P. grisea* displays deep purple nectar guides typical of most *Physalis*, whereas *P. pruinosa* does not. (**Figure 2A**). This pigmentation difference is also found on stems and branches. Crossing *P. grisea* and *P. pruinosa* resulted in F1 hybrids showing purple pigmentation, and an F2 population showed that yellow color segregated as a single recessive mutation. Mapping-by-sequencing localized the mutation to chromosome 4; however, limited recombination resulted in a large interval spanning most of the chromosome (**Figure 2B**).

**Figure 2.**
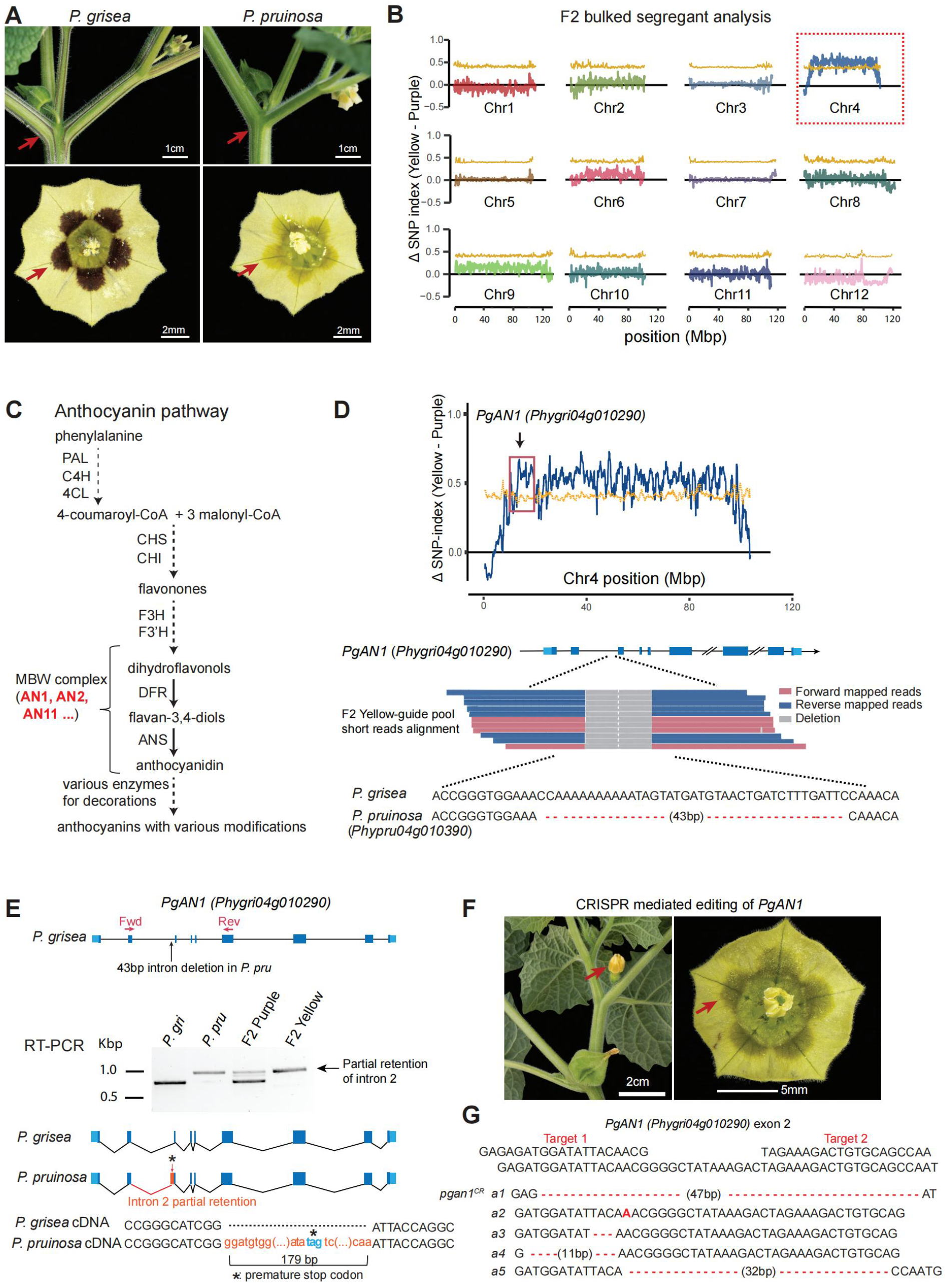
Loss of purple pigmentation in *P. pruinosa* is due to an intronic SV in the bHLH transcription factor gene *ANTHOCYANIN1*. **A.** Images showing difference in pigmentation between *P. grisea* and *P. pruinosa*. Arrows point to purple (*P. grisea*) compared to yellow (*P. pruinosa*) pigmentation on stems and flowers. **B.** Mapping-by-sequencing showing ΔSNP-index across all 12 chromosomes using *P. grisea* as the reference, with SNP ratios between yellow-guide and the purple-guide pools from an inter-specific F2 population. Yellow line: 95% confidence interval cut-offs of ΔSNP-index. **C.** Simplified pathway of anthocyanin biosynthesis based on data from petunia. Major transcriptional and enzymatic regulators are shown as abbreviations. Dashed lines indicate multiple steps condensed. Bold red font indicates components of the MYB-bHLH-WD40 (MBW) complex that transcriptionally activates late biosynthetic genes. **D.** Top: The ΔSNP-index plot for Chromosome 4. Black arrow points to the genomic location of the *AN1* candidate gene. Bottom: pile-up of Illumina mapped-reads from *P. pruinosa* at the 2^nd^ intron of *AN1* showing a 43-bp deletion in *PpAN1* (*Phypru04g010390*) all sequences. In all gene models (including later figures), deep blue boxes, black lines, and light blue boxes represent exonic, intronic, and untranslated regions, respectively. **E.** Molecular consequences of the 43-bp intronic deletion in *PpAN1* revealed by RT-PCR and sequencing. Red arrows indicate RT-PCR primers. Longer amplicons and thus *AN1* transcripts from both the yellow-guide F2 bulk pool and *P. pruinosa* were identified by agarose gel electrophoresis. Sanger sequencing revealed the inclusion of a 179 bp fragment of intron 2 in the *PpAN1* amplicon, resulting in a premature stop codon. Red box reflects intronic sequence retained in the transcript. Black asterisk, premature stop. **F.** Loss of purple pigmentation in CRISPR edited *PgAN1* T_0_ plants. **G.** CRISPR-Cas9 generated mutant alleles from the yellow T_0_ chimeric plants are shown. Red dashed lines represent deletions. Red bold letter indicates a single nucleotide insertion.

To identify candidate genes, we searched for homologs of genes involved in the production of anthocyanins in the *Solanaceae* genus *Petunia*. Anthocyanins belong to a class of polyphenolic secondary metabolites named flavonoids, and one outcome of their accumulation in tissues and organs is purple pigmentation (Liu et al., 2018) . Many ornamental *Petunia* species show variation in anthocyanin accumulation, and studies on this diversity have identified enzymes and transcription factors in the anthocyanin pathway (Bombarely et al., 2016; Liu et al., 2018).

Anthocyanin biosynthesis involves three major steps, including the conversion of phenylalanine to 4-coumaroyl-CoA through stepwise enzymatic reactions, and the conversion of 4-coumaroyl-CoA to dihydroflavonols, which are precursors in the final synthesis steps of specific anthocyanins (**Figure 2C**). We identified four orthologs of anthocyanin pathway genes on chromosome 4. Overlaying our SV analysis revealed a mutation in only one of these genes, a 43 bp deletion in the second intron of the *P. pruinosa* gene *Phypru04g010390*, which encodes a bHLH transcription factor orthologue of petunia ANTHOCYANIN1 (AN1) (Spelt et al., 2000) (**Figure 2D**). AN1 activates the structural gene *DIHYDROFLAVONOL REDUCTASE* and other anthocyanin regulators (Spelt et al., 2000) . Notably, mutations in petunia *AN1* result in loss of anthocyanins in all tissues (Spelt et al., 2000, 2002). Using qRT-PCR and sequencing of cDNA we found that *AN1* transcripts in *P. pruinosa* were longer than those in *P. grisea* due to a retention of 179 bp from intron 2, which results in a premature stop codon (**Figure 2E**). We validated this result by CRISPR-Cas9 targeting *PgAN1* (*Phygri04g010290*) in *P. grisea*. Five out of 11 first generation (T_0_) transgenic lines failed to produce anthocyanins and sequencing showed these plants carried edited alleles of *PgAN1* (**Figure 2F, G**). These results indicate that the SV in *P. pruinosa AN1* (*PprAN1*) underlies the loss of pigmentation in *P. pruinosa* and further demonstrate the utility of our genomic resources in deploying forward genetics in *Physalis*.

### The MADS-box genes *MPF2* and *MPF3* are not essential regulators of ICS

The most striking feature of *Physalis* is the inflated calyx syndrome (ICS), which evolved repeatedly in other *Solanaceae* genera and angiosperms (Deanna et al., 2019; Padmaja et al., 2014; Paton, 1990) . Soon after fertilization, sepals undergo remarkable growth and expansion acropetally to encapsulate fruits in balloon-like papery husks, which may provide protection from pathogens and promote seed dispersal (**Figure 3A**) (Baumann & Meier, 1993; J. Li et al., 2019) . Despite long-standing interest, the evolutionary and mechanistic origins of ICS remain unclear. One early defining study proposed that heterotopic expression of *MPF2* was essential to the evolution of ICS (He & Saedler, 2005) . This hypothesis was based on over-expression of the potato ortholog, *StMADS16,* in the tobacco species *N. tabacum*, which produced leaf-like sepals. Empirical support in *Physalis* came from RNA interference (RNAi) knock-down of *MPF2* in *P. floridana,* where multiple transgenic lines showed a reduced calyx size, the severity of which was highly correlated with impaired fertility, but counter-intuitively not the level of reduction of *MPF2* transcripts (He & Saedler, 2005).

**Figure 3.**
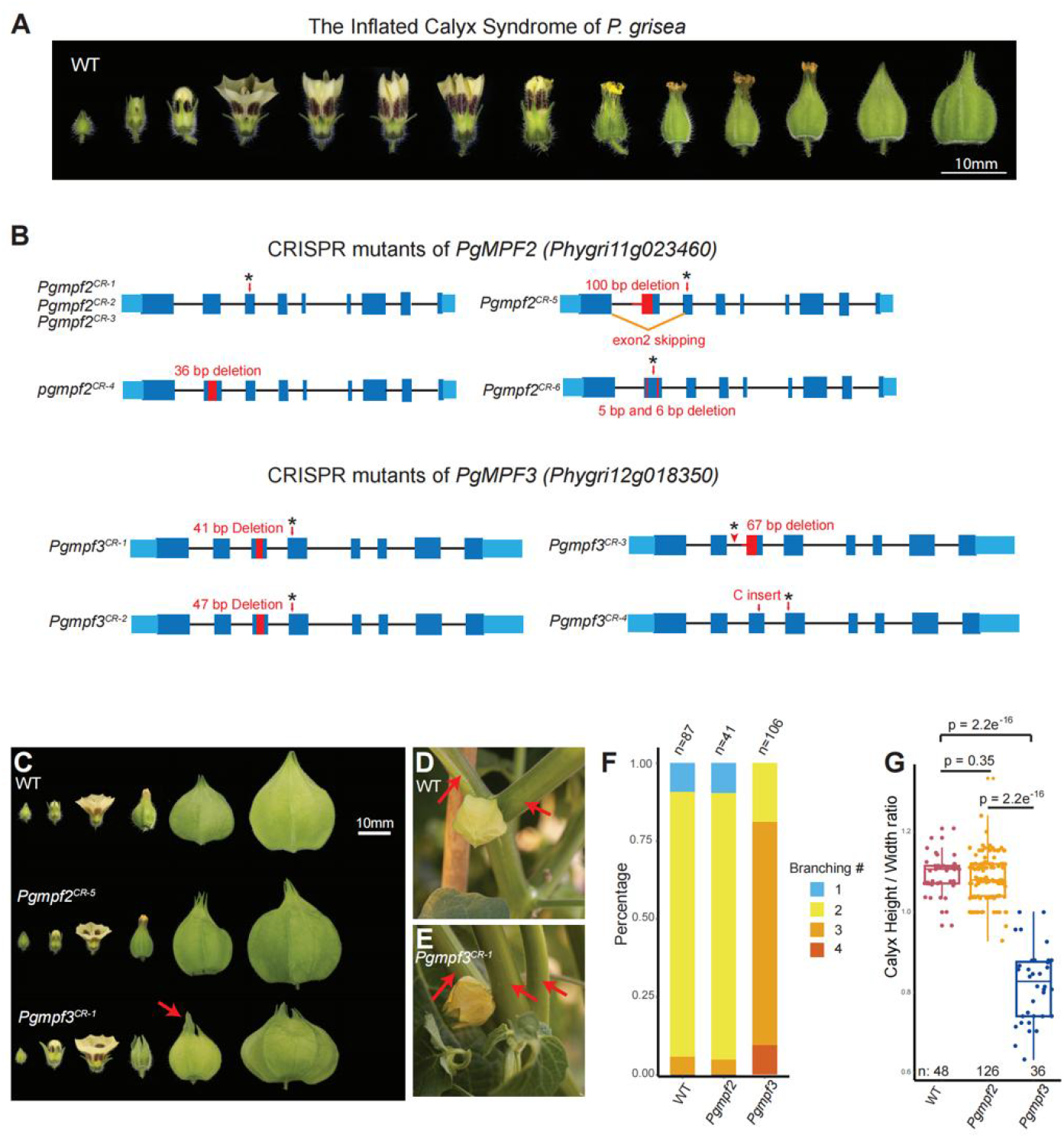
CRISPR-Cas9 generated mutants of the MADS-box genes *PgMPF2* and *PgMPF3* do not prevent ICS. **A.** Images showing sequential stages of ICS in *P. grisea* from early flower formation to calyx inflation over 3 days. **B.** Multiple, independently-derived null alleles in *PgMPF2* and *PgMPF3*. Red boxes and lines, deletions; black asterisks, stop codons. Three alleles of *PgMPF2* (*Pgmpf2^CR-1^*, *Pgmpf2^CR-2^*, *Pgmpf2^CR-3^*) with different mutations in exon 3 result in the same premature stop codon. Specific mutations for all alleles are shown in **Supplemental Figure. S2** and in **Source Data 1. C-G.** Phenotypes of *Pgmpf2* and *Pgmpf3* null mutants. All homozygous mutants independently derived alleles showed the same phenotypes, and *Pgmpf2^CR-5^* and *Pgmpf3^CR-1^* were used as references for phenotypic analyses. **C.** Calyx inflation is not disrupted in *Pgmpf2* and *Pgmpf3* mutants. Representative images from *Pgmpf2^CR-5^* and *Pgmpf3^CR-1^* are shown. The leaf-like sepal tip of *Pgmpf3^CR-1^* is indicated by the red arrow. **D and E.** Shoot branching phenotype of *Pgmpf3^CR-1^* compared to WT. A typical sympodial unit of WT *Physalis* consists of one leaf, one flower and two side shoots. *Pgmpf3* mutants develop mostly three side shoots (**E**). Branches are indicated by red arrows in representative images. **F.** Quantification of branching in WT, *Pgmpf2* and *Pgmpf3* shown as stacked bar charts. **G.** Quantification of calyx height/width ratio in WT, *Pgmpf2* and *Pgmpf3*. Statistical significance determined by two-tailed, two-sample *t*-tests, and *p* values are shown.

Despite this contradictory result, follow-up studies proposed and tested an extended mechanism involving regulation of *MPF2* by the *AP1*-like transcription factor gene *MPF3* (orthologue of *Arabidopsis AP1: APETALA1* and tomato *MC: MACROCALYX)*, in combination with hormonal control and fertilization (He & Saedler, 2007; Zhao et al., 2013) . However, functional data supporting these conclusions were based on over-expression and also RNAi and Virus Induced Gene Silencing (VIGS) knockdown of expression. Pleiotropic phenotypic outcomes are common in over-expression experiments, and challenging to relate to specific genes studied, whereas RNAi and VIGS are difficult to interpret due to variable knock-down efficiencies and potential off-target effects (Senthil-Kumar & Mysore, 2011; P. Xu et al., 2006) . Further convolution of a possible ICS mechanism emerged with the recent publication of the *P. floridana* genome, and the suggestion that absence of the *SEP4* orthologue of the tomato MADS-box gene *SlMBP21*/*J2* in *Physalis* was yet another critical factor in the origin of ICS (Lu et al., 2021).

To address these inconsistencies and provide a more robust genetic dissection of ICS, we first used CRISPR-Cas9 genome editing to eliminate *MPF2* and *MPF3* function in *P. grisea*. We generated five alleles of *PgMPF2 (Phygri11g023460)* and four alleles of *PgMPF3 (Phygri12g018350)* (**Figure 3B**), and these independent mutations caused different premature stop codons. Notably, none of these homozygous mutants disrupted ICS; all *Pgmpf2^CR^* mutants showed similar calyx inflation as WT, and *Pgmpf3^CR^* mutants displayed enlarged and more leaf-like tips of sepals before inflation, a phenotype also observed in tomato *mc* mutants (**Figure 3C**) (Yuste-Lisbona et al., 2016) . While this change of sepal tips is accompanied by a lower calyx height/width ratio (**Figure 3G**), inflation was unaffected. Besides the sepal phenotype, *Pgmpf3* also displayed abnormal branching patterns; *Pgmpf3* mutants frequently produce three instead of two sympodial shoots (**Figure 3D-F**). In summary, these CRISPR-Cas9 engineered loss-of-function mutations in *PgMPF2* and *PgMPF3* show that these MADS-box genes alone are not responsible for the evolution of ICS and are not essential regulators of this developmental process.

### Targeted mutagenesis of additional MADS-box genes does not abolish ICS

In an effort to identify genes involved in ICS, we embarked on a more comprehensive reverse genetics approach targeting MADS-box genes known to regulate floral organ development in tomato and other species, including additional MADS-box family members that mimic ICS when over-expressed or mutated in non-ICS *Solanaceae*. For example, we characterized a spontaneous tomato mutant with greatly enlarged fleshy fruit-covering sepals and found a transposon insertion SV upstream of *TOMATO AGAMOUS-LIKE1* (*TAGL1*), that caused >80-fold over-expression in developing sepals (**Figure 4A**). *TAGL1* belongs to the *AGAMOUS* clade of MADS-box transcription factors, and is a close paralog of *TOMATO AGAMOUS 1 (TAG1).* Previous studies showed both of these genes control flower development, and when either is over-expressed, enlarged and fleshy sepals are produced, in part mimicking ICS (Itkin et al., 2009; Pnueli et al., 1994). To test the roles of the *Physalis* orthologues of these genes, we generated CRISPR mutants. As observed in corresponding mutants of other species (Pan et al., 2010; Yanofsky et al., 1990), *Pgtag1^CR-1^* homozygous mutants displayed severe homeotic transformation of stamens to petal-like structures, while *Pgtagl1^CR-1^* displayed similar but weaker homeotic transformations (**Figure 4B**). Importantly, despite these floral organ defects, accompanied also by partial or complete loss of self-fertilization, both of these mutants maintained inflation, although calyx size was reduced, potentially due to secondary growth effects (**Figure 4B-E**).

**Figure 4.**
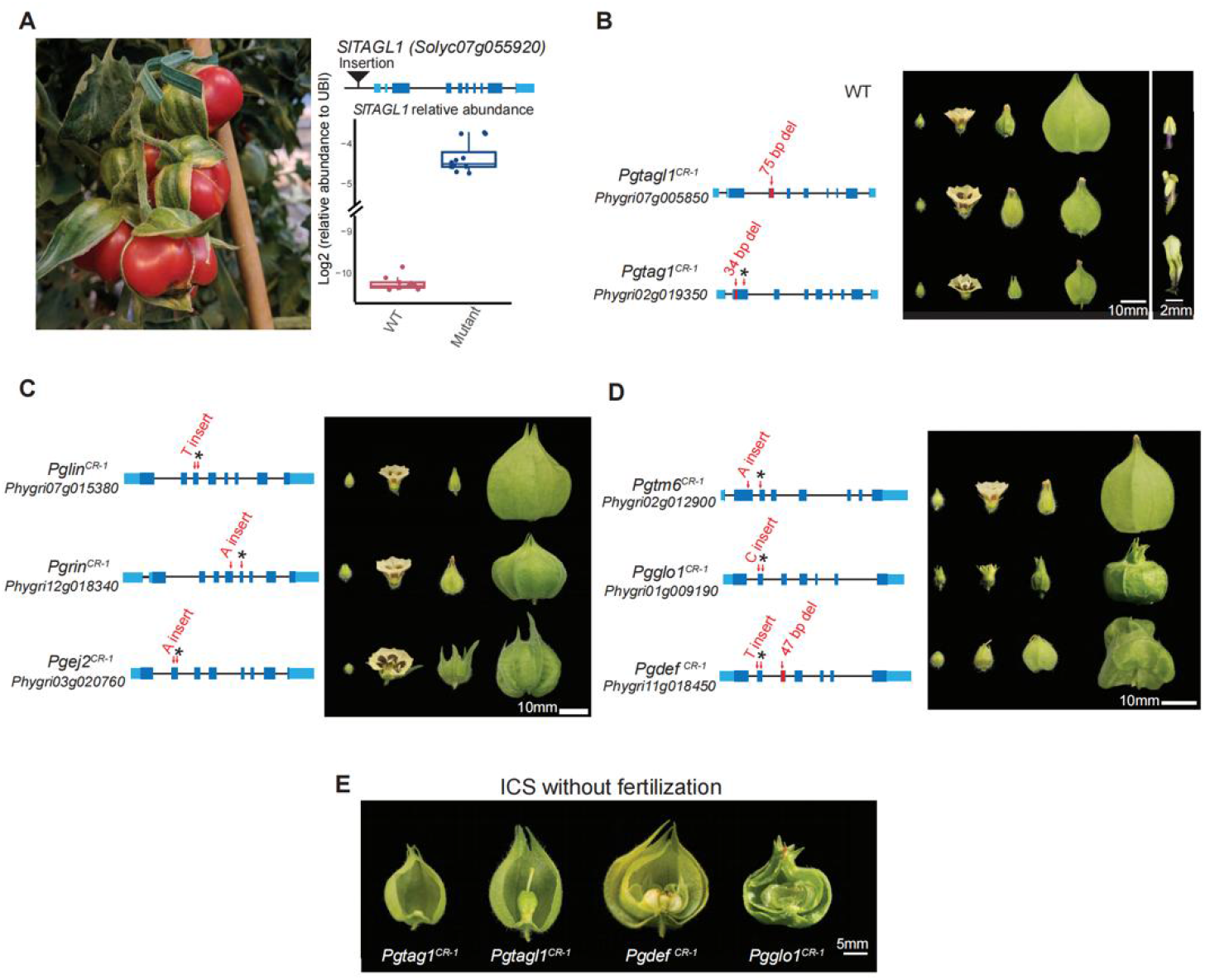
CRISPR-Cas9 generated mutations in eight additional candidate MADS-box genes do not disrupt ICS. **A.** Over-expression of *SlTAGL1* caused by a transposable element insertion (see **Methods**) results in enlarged calyx in tomato, mimicking ICS and presenting another candidate MADS-box gene. Left: image of calyx phenotype from the *SlTAGL1* mutant. Right, top: gene model of *SlTAGL1* with the transposon insertion (black triangle) identified by genome-sequencing. Right, bottom: qRT-PCR on cDNA derived from young sepals showing over-expression of *SlTAGL1* in the mutant. Three biological replicates for WT, and four biological replicates for the mutant were assayed; each data point represents one technical replicate. **B.** Mutations in *PgTAGL1* and *PgTAG1* cause homeotic transformations of stamens to petal-like organs but do not disrupt ICS. Middle image: representative calyx phenotypes at different developmental stages. Right image: single organs from the 3^rd^ floral whorl. **C.** Mutations in three *SEP4* homologs do not disrupt ICS. **D.** Mutations in multiple B-function MADS-box genes do not disrupt ICS. **E.** ICS still occurs in mutants with fertilization defects or that fail to produce fruits. Mutations in *PgTAG1*, *PgTAGL1*, *PgDEF*, and *PgGlo1* cause homeotic transformations of floral organs that abolish self-fertilization, but ICS is preserved.

Based on their roles in floral organ development and inflorescence architecture, *SEPALLATA4 (SEP4*) MADS-box genes are another set of ICS candidates. Tomato has four *SEP4* clade MADS-box genes: *J2*, *SlMADS1/ENHANCER OF J2* (hereafter *EJ2*), *LONG INFLORESCENCE* (*LIN*) and *RIPENING INHIBITOR* (*RIN*). We previously showed that *EJ2* and *LIN* regulate sepal development; mutants of *ej2* alone and in combination with *lin* develop enlarged sepals (Soyk et al., 2017). Analysis of the genome of *P. floridana* (Lu et al., 2021), and confirmed in our genomes, showed that *Physalis* lost the ortholog of *J2*, whereas the other three *SEP4* genes are present. Curiously, loss of *J2* was proposed to have promoted the evolution of ICS, but non-ICS *Solanaceae* such as pepper also lack *J2*. To test roles of the *SEP4* clade in ICS, we used CRISPR-Cas9 to mutate all three *SEP4* genes in *P. grisea*. Notably, multiple independent mutations in *PgEJ2, PgLIN,* and *PgRIN* did not inhibit ICS. Similar to our findings in tomato *ej2* mutants (Soyk et al., 2017), mutants of *Pgej2^CR-1^* produced larger sepals in young and fully developed flowers, but inflation proceeded normally, with the only modification being sepal tips fail to coalesce to a single point after inflation is complete (**Figure 4C**).

### Fertilization is not required for ICS

In flower development, B-class MADS-box genes participate in specifying petal and stamen identity, and the loss of B function leads to homeotic transformations of petals and stamens, which impair self-fertilization (Theißen & Saedler, 2001; Weigel & Meyerowitz, 1994; Yanofsky et al., 1990). If fertilization-related signals were required for ICS, as reported (He & Saedler, 2007), mutations in B-class MADS-box genes should result in abnormal ICS development. Previously, a mutation deleting the B-class MADS-box gene *GLOBOSA1* (*GLO1)* was shown to develop a double-layered calyx phenotype in *P. floridana* when fertilized with WT pollen (J.-S. Zhang et al., 2014) . We identified four B-class MADS-box genes in *P. grisea*, including the four closest homologs of *GLO1*: *PgGLO1* (*Phygri01g009190*), *PgGLO2* (*Phygri06g017940*), *PgDEF* (*Phygri11g018450*) and *PgTM6* (*Phygri02g012900*). CRISPR-Cas9 induced null mutations in all four genes failed to disrupt ICS. Mutants of *Pgtm6^CR-1^* and *Pgglo2^CR-1^* appeared WT, whereas *Pgglo1^CR-1^* and *Pgdef^CR-1^* both displayed expected homeotic transformations of stamens to carpels, and petals to sepals. Notably, calyx inflation was unaffected even in the second whorls of *Pgglo1^CR-1^* and *Pgdef^CR-1^* where petals were converted to sepals (**Figure 4D, E**).

Fertility or signals from developing fruits have also been observed to be required for the initiation and progression of inflation, perhaps due to the activity and signaling of hormones such as cytokinin and gibberellin (He & Saedler, 2007) . However, many of our MADS-box mutants with severe floral organ homeotic transformations also fail to self-fertilize, and have various degrees of defects in fruit development. That ICS is unaffected in these mutants provides compelling genetic evidence that ICS can be uncoupled from normal fertilization. In particular, both *Pgdef ^CR-1^* and *Pgglo1^CR-1^* homozygous mutants cannot self-fertilize and form multiple small fruits without seeds due to homeotic transformations of stamens to carpels, yet the twin outer layers of sepals still form inflated calyces (**Figure 4E**). Moreover, in *Pgtagl1^CR-1^*and *Pgtag1^CR-1^*mutants, which cannot self-fertilize and whose fruits arrest early in development or fail to form entirely, respectively, inflation remained intact (**Figure 4E**).

In summary, although earlier observations, hypotheses, and data suggested critical roles of several MADS-box genes in the evolution of ICS, our results show that calyx inflation is maintained in loss-of-function mutants of the *P. grisea AG* clade, *SEP4* clade and B-class MADS-box transcription factor genes. These data further demonstrate that although fertilization signals or developing fruit may contribute to the regulation of calyx inflation, neither is absolutely required.

### *Huskless*, a mutation in an *AP2*-like transcription factor, eliminates inflated calyx

Forward genetics is a powerful and unbiased approach to identify genes controlling traits of interest in model systems. We performed a small-scale ethyl methanesulfonate (EMS) mutagenesis screen in *P. grisea* to identify genes involved in calyx development (**see Methods**). A recessive mutant bearing fruits without husks was identified and named *huskless* (*hu*) (**Figure 5A, B**). Scanning electron microscope (SEM) imaging of dissected flower buds showed that *hu* mutants developed three floral whorls instead of four compared to WT (**Figure 5C, D**). To isolate the causative mutation, we sequenced genomic DNA from a pool of *hu* mutants and WT siblings from the original *P. grisea* mutagenesis (M2) family (**see Methods**). Aligning Illumina-sequenced reads to the *P. grisea* genome allowed screening for single nucleotide variants (SNVs) that were homozygous in the *hu* pool but not in the WT sibling pool. We scored these SNVs for predicted functional consequences on annotated gene transcripts using SnpEff (Cingolani et al., 2012) . Out of eight such SNVs, one was a G-to-A mutation in a 3’ splice site of *Phygri09g010120*, which encodes an *APETALA2* (*AP2*)-like transcription factor (**Figure 5E; Supplemental Table S6a**). Co-segregation analysis in M3 families confirmed association of this mutation with the *hu* phenotype (**Supplemental Table S6b**), and sequencing RT-PCR products of *Phygri09g010120* from *hu* floral tissue showed mis-splicing in the 4^th^ intron, resulting in partial skipping of exon 5 (**Figure 5E**). Importantly, independent CRISPR generated mutations of this *AP2-like* gene in *P. grisea* resulted in independent mutations that caused the same phenotype as *hu* **(Figure 5F)**.

**Figure 5.**
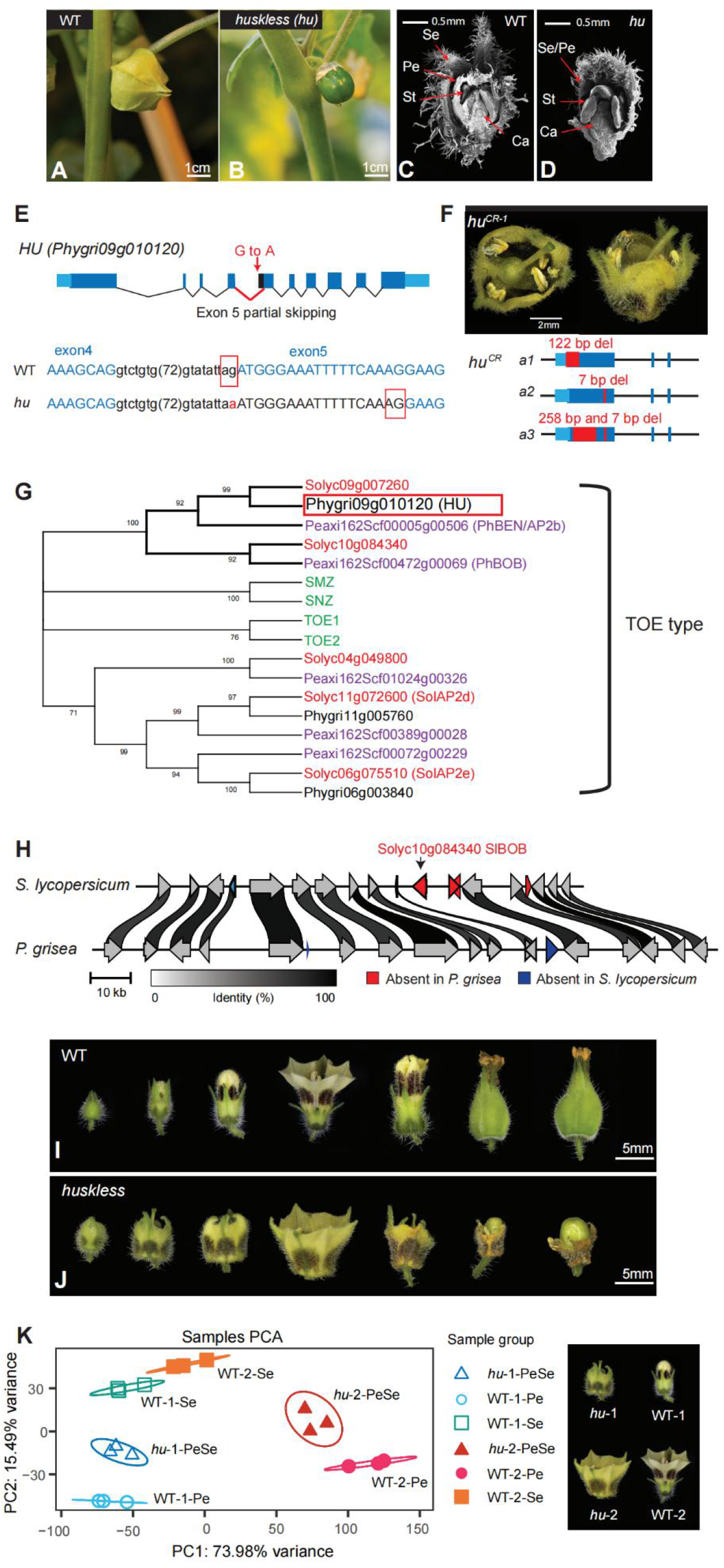
The *huskless* mutant lacks an inflated calyx due to mutation of an *AP2*-like transcription factor. A-D. Phenotypes of the EMS-derived *huskless* (*hu*) mutant. **A and B.** Images of WT and *hu* displaying the loss of calyx phenotype at the mature green fruit stage. **C and D.** Longitudinal SEM images of WT and *hu* developing flowers showing *hu* mutants develop only three floral whorls compared to four in WT. The first whorl of *hu* flowers shows hallmarks of sepal and petal identity. Se: sepal; Pe: petal; St: stamen; Ca: Carpel. **E.** Gene model showing the G-to-A point mutation causing partial skipping of exon 5 in the *AP2-like* transcription factor gene *Phygri09g010120.* Blue colored nucleotides represent exonic sequences; red boxes indicate 3’ splice sites in WT and *hu*. **F.** CRISPR-Cas9 generated mutations in *Phygri09g010120.* Top: gene models showing three independent CRISPR null alleles of *hu*. Sequences 3’ of the 3^rd^ intron are omitted. *hu^CR-1^* is homozygous for allele 1 (*a1*). Bottom: images of *hu^CR-1^* flower phenotype. **G.** Maximum Likelihood consensus tree of the TOE-type euAP2 proteins from *A. thaliana* (gene names in green), *P. axillaris* (Peaxi IDs in purple), *S. lycopersicum* (Solyc IDs in red), and *P. grisea* (Phygri IDs in black). Bootstrap values (%) based on 500 replicates are indicated near the branching points; branches below 50% have been collapsed. **H.** Local synteny analysis between *S. lycopersicum* and *P. grisea* showing the absence of the *Solyc10g084340* orthologue (petunia *BOB* orthologue) in *P. grisea*. Arrows indicate genes and orientations. Protein identity percentages between orthologues are indicated by ribbon shades in grayscale; only links above 80% identity are shown. **I and J.** Series of images of WT and *hu* developing flowers from before anthesis through early fruit development. **K.** Principal component analysis (PCA) of WT and *hu* RNA-seq data. Right image: visual reference of the two stages used for expression profiling from WT and *hu* floral whorls. Numbers (-1 or -2) in the sample groups represent stage 1 or 2; petal or sepal whorls in WT are denoted as Pe, Se respectively; PeSe represents the merged outer whorl in *hu*. The top 3000 differentially expressed genes were used for PCA.

*HU* is the homolog of *Petunia hybrida AP2B/BLIND ENHANCER* (*BEN*) (**Figure 5G**), which specifies 2^rd^ and 3^rd^ floral whorl identity (Morel et al., 2017) with its redundant paralog *BROTHER OF BEN* (*BOB*). Petal development is strongly inhibited in *ben bob* double mutants, resulting in severely reduced or absent petals, and partial conversion of sepals into petals, resembling *hu* (Morel et al., 2017) . Because the *Petunia hybrida* genome is highly fragmented (Bombarely et al., 2016), we performed a synteny analysis of the chromosomal segments containing *BOB* in *P. grisea, P. pruinosa*, and *S. lycopersicum* and found that this paralog of *HU* (*BEN*) is present in tomato but not in groundcherry (**Figure 5H**). Thus, *hu* emerged in our forward genetics mutagenesis screen, because the *BOB* ortholog and therefore redundancy is absent in *P. grisea*.

The first floral whorl of *hu* displays characteristics of both sepals and petals (**Figure 5I, J**). The whorl begins developing with green as the dominant color, like sepals, but gradually turns yellow as the flower matures, maintaining green color at organ tips. Nectar guides are also visible throughout development of the first whorl, indicative of early petal identity. After fertilization, the first whorl mildly increases in size but fails to fully inflate before gradually senescing as *hu* fruits develop into the size of WT fruits.

To characterize the role of *HU* in whorl identity and ICS, we profiled transcriptomes by RNA-seq from WT sepals and petals at two stages of organ maturation and compared them with corresponding stages of *hu* first whorls (**Figure 5K, and Methods**). Principal component analysis (PCA) revealed *hu* expression profiles (denoted as *hu*-PeSe) were positioned between the profiles of WT sepals and petals at both stages, supporting the mixed organ identity observed phenotypically. Thus, the loss of the inflated calyx in *hu* mutants is from a failure to properly specify sepal and petal identity as opposed to directly disrupting a mechanistic origin of ICS. Our findings also demonstrate how lineage-specific gene duplications and paralog presence-absence variation can shape genetic redundancies and genotype-to-phenotype relationships over short time frames, and further illustrate the value of multiple related model systems.

## DISCUSSION

Discoveries in plant development, cell biology, and genetics continue to depend on a limited number of model systems, historically centered around *Arabidopsis thaliana* and its relatives in the Brassicaceae family (Chang et al., 2016). New models are essential to advance fundamental and applied research beyond the small fraction of biological diversity captured by current models. While additional model species have been proposed or are under development (Chang et al., 2016), most lack the powerful combination of efficient genomics and genetics. Moreover, emphasis is largely on neglected lineages and single representative species within them. An approach with complementary benefits relies on multiple models within a lineage to address often overlooked questions of species-specific and comparative evolutionary history over short time frames. The *Solanaceae* family is ideal in this regard, including i) rich diversity throughout ∼100 genera and more than 3000 species spanning ∼30 million years of evolution, ii) broad agricultural importance from more than two dozen major and minor fruit and vegetable crops, and iii) feasibility of rapidly developing and integrating genome editing with reference and pan-genome resources.

By establishing high-quality chromosome-scale assemblies for *P. grisea* and *P. pruinosa,* we cemented these *Physalis* as new models and advanced *Solanaceae* systems with genomics and genetics. Most significantly, our integration of these resources revealed that the mechanisms underlying ICS remain elusive. Indeed, despite previous evidence suggesting otherwise, we conclude that none of the 11 candidate MADS-box genes we functionally characterized using genome editing, nor fertility alone, are core regulators of ICS. Our findings therefore force a reset in the search for the physiological, genetic, and molecular mechanistic origins of this evolutionary novelty. Though a logical starting point, the candidate gene approach based on MADS-box over-expression phenotypes in other species was prone to misleading hypotheses and false positives, likely due to the complex evolutionary history of the MADS-box family members and their even more complex genetic and physical interactions. Indeed, multiple MADS-box genes appear to be capable of mimicking ICS through over-expression, possibly due to coordinated activation of closely related paralogs and subsequent complex feedback regulation and interactions among other family members. This might suggest double and higher order mutants of these or other MADS-box genes not investigated here would ultimately perturb ICS, possibly reflecting a collective role of multiple family members acting redundantly or in a network. However, such a result would not necessarily indicate direct roles for these genes in the evolutionary steps leading to ICS.

Based on our genetics, we expect additional or other genes and molecular programs are central, and the tools established here provide the foundation to revisit ICS in an unbiased way. ICS is a rapid and dynamic process, where extraordinary morphological changes in sepal growth and inflation occur within a few days. This suggests that the molecular events driving and responding to the inception of the transition from a non-inflated sepal whorl to active inflation may be short-lived, happening in the order of hours. We propose that the future dissection of ICS should be based on detailed and integrated temporal, morphological and molecular analyses to capture these transient events. A recent study in tomato took advantage of transcriptome profiling and computational ordering of hundreds of single shoot apical meristems to capture and reconstruct a highly detailed temporal gene expression map of the floral transition. These data revealed previously hidden gene, short-lived expression programs and several genes that function in parallel transient pathways critical to the floral transition process (Meir et al., 2021). With the new reference genomes and annotations of *P. grisea* and *P. pruinosa*, a similar approach can be applied to ICS, where large numbers of individual sepals can readily and reliably be harvested and profiled throughout calyx development. As opposed to focusing on entire floral buds (H. Gao et al., 2020), such high-resolution temporal transcriptome profiling of sepals alone would provide comprehensive and unbiased information regarding global and possibly gene-specific molecular signatures in the initiation and maintenance of inflation, and expose new candidates that can be studied using the integrated genomics and genome editing strategies demonstrated here.

Beyond floral development and ICS in *Physalis*, our work sets a high-quality anchor to broaden biological questions and discoveries in the *Solanaceae*, and further illustrates fast and efficient approaches to building new model systems. Establishing new pan-genome and genome editing tools in many additional genera of *Solanaceae* and of other plant families will enable comparative genomic and genetic studies over both short and long evolutionary timescales.

---

**Supplemental Figure S1.**
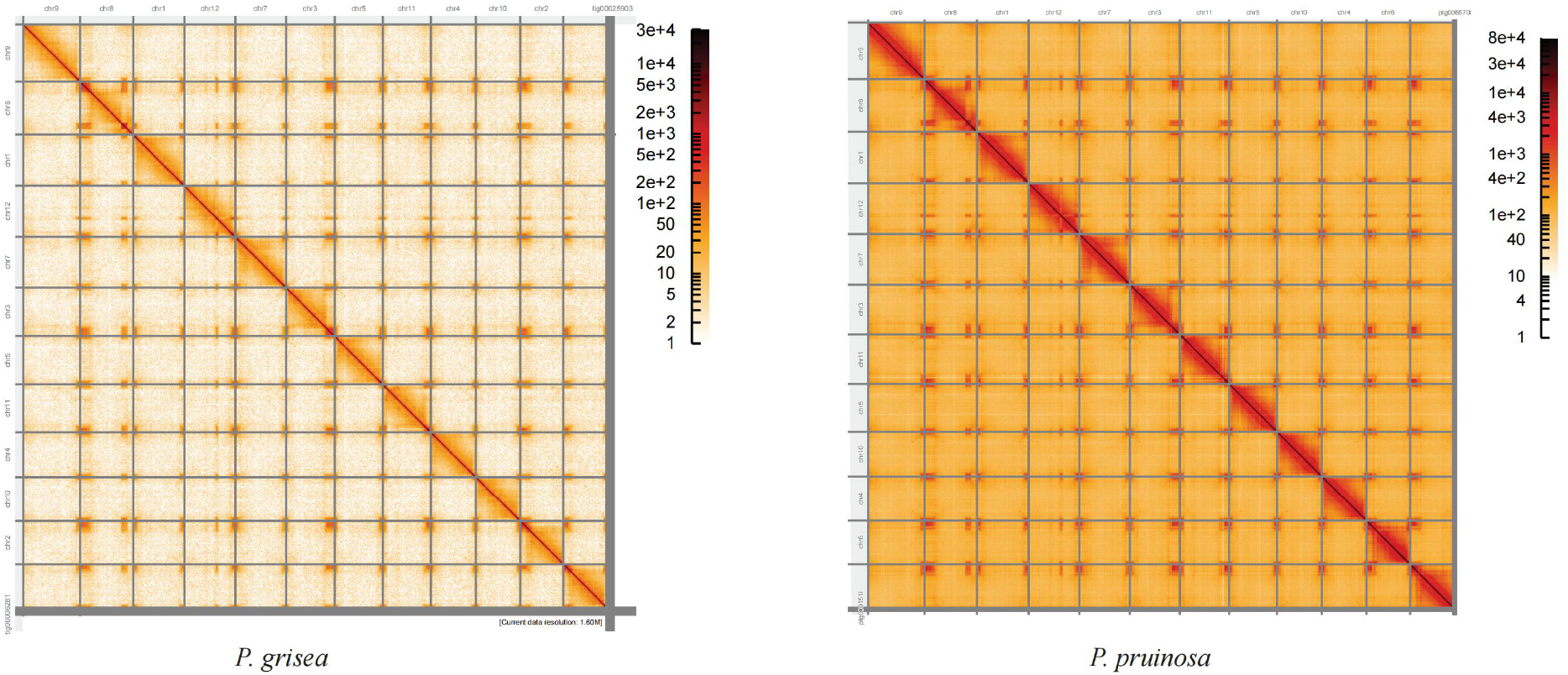
Hi-C heatmaps confirm reference assembly structural accuracy. Hi-C heatmaps for the *P. grisea* and *P. pruinosa* reference assemblies. The 12 chromosomes are sorted from largest (top left) to smallest (bottom right).

**Supplemental Figure S2.**
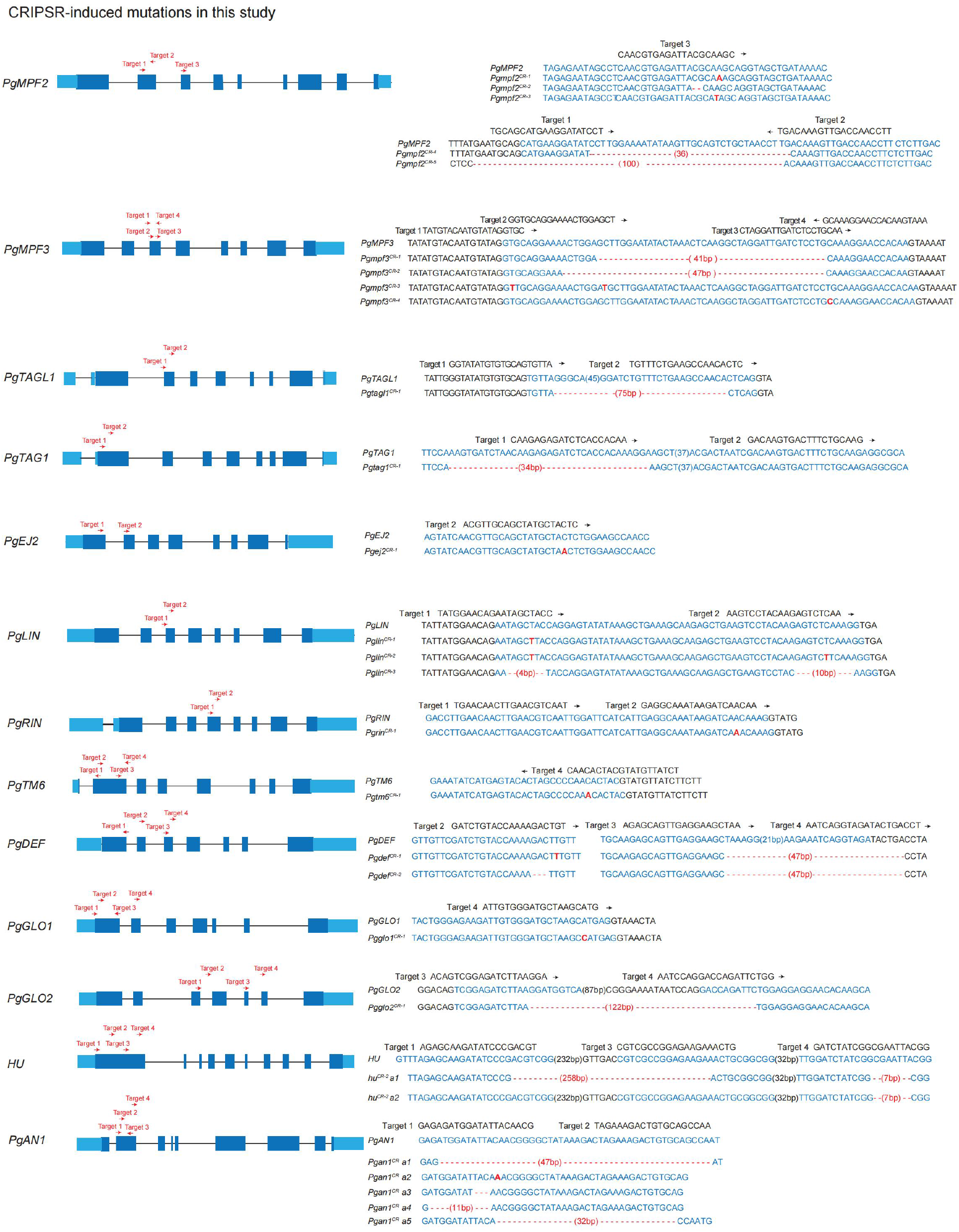
Illustrations of CRISPR-engineered mutations in this study. In all gene models, deep blue boxes, black lines, and light blue boxes represent exonic, intronic, and untranslated regions, respectively. Red dashed lines indicate indels/deletions; Red bold nucleotides represent inserts.

**Supplemental Figure S3.**
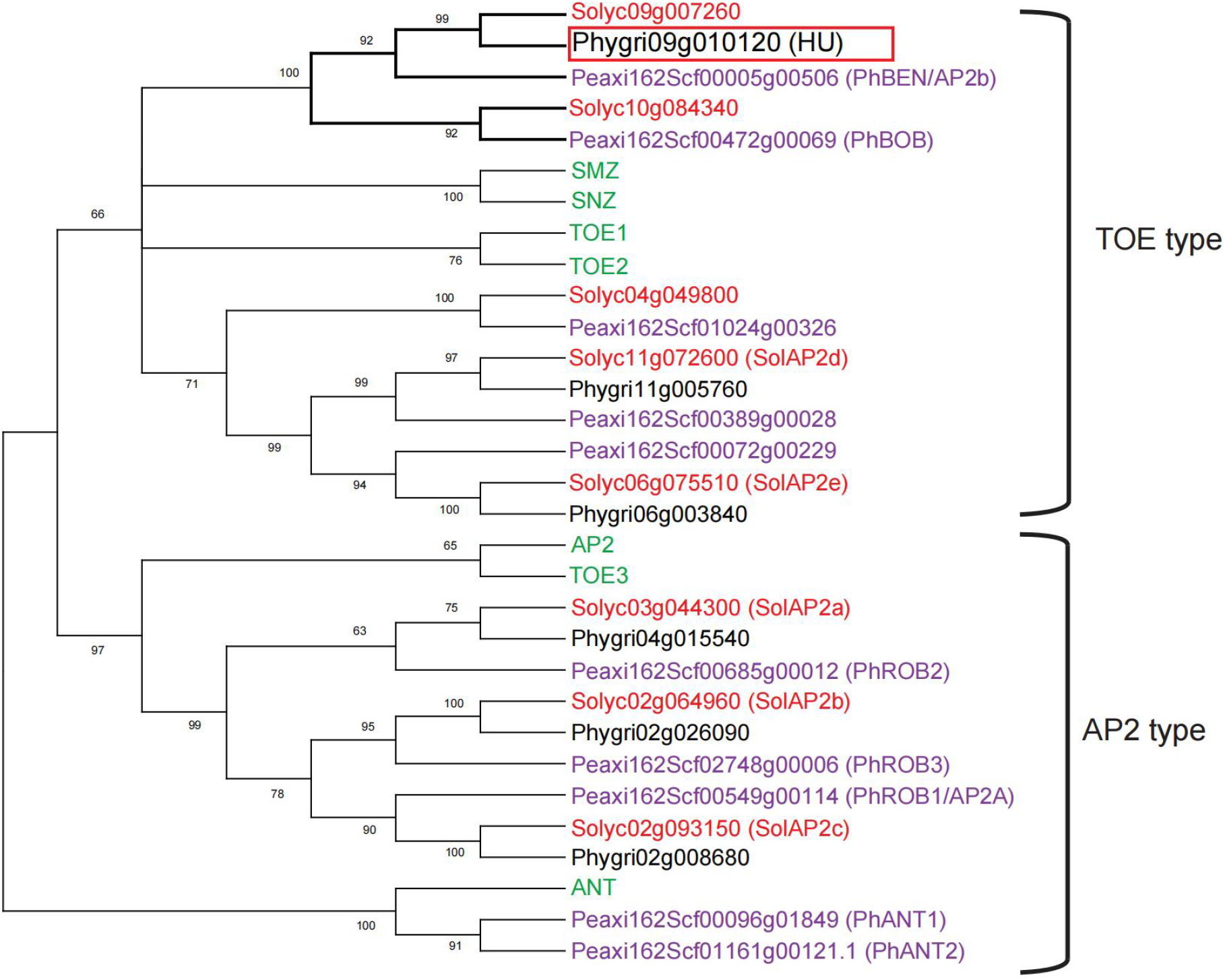
Maximum Likelihood consensus tree of the euAP2 proteins from *A. thaliana* (gene names in green), *P. axillaris* (Peaxi IDs in purple), *S. lycopersicum* (Solyc IDs in red), and *P. grisea* (Phygri IDs in black). Bootstrap values (%) based on 500 replicates are indicated near the branching points; branches below 50% have been collapsed. Protein sequences used to build the tree can be found in **Source Data 6**.

## Materials and methods

### Plant material, growth conditions, and phenotyping

Seeds of *P. grisea* and *P. pruinosa* were obtained from the Solanaceae Germplasm Bank at the Botanical Garden of Nijmegen and from commercial seed sources. Seeds were directly sown into soil in 96-well plastic flats and grown in the greenhouse under long-day conditions (16-hr light/8-hr dark) supplemented with artificial light from high-pressure sodium bulbs (∼250 μmol m^−2^ s^−1^). The temperature ranged from 26-28°C during the day to 18-20°C during the night, with a relative humidity of 40%–60%. 4-week old seedlings were transplanted to 4 L pots in the same greenhouse, or into the fields at Cold Spring Harbor Laboratory (CSHL) unless otherwise noted. The tomato mutant displaying enlarged fleshy sepals from **Figure 4** was a gift from Dr. Dani Zamir, which arose from the whole genome backcross lines constructed from a cross between *Solanum pimpinellifolium* (LA1589) and *Solanum lycopersicum* inbred variety cv. E6203 (TA209) (Grandillo & Tanksley, 1996). Branching and internode length phenotypes were assayed in greenhouse grown plants at 2 months after sowing.

### Extraction of high-molecular weight DNA and long-read sequencing

For long-read sequencing, shoot apices of 3-week old seedlings were harvested after a 48-hr dark treatment. Extraction of high-molecular weight genomic DNA, construction of Oxford Nanopore Technology (ONT) libraries and PacBio HiFi libraries, and sequencing were described previously (Alonge et al., 2020, 2021) . Hi-C experiments were conducted at Arima Genomics (San Diego, CA) from 2 g of flash-frozen leaf tissue.

### *P. grisea* chloroplast and mitochondria genome assembly

To assemble the *P. grisea* chloroplast genome, all HiFi reads were aligned to the previously published *Physalis* chloroplast reference genome (GenBank ID MH019243.1) with Minimap2 (v2.17-r974-dirty, -k19 -w19) (H. Li, 2018) . All reads with at least one primary alignment spanning at least 90% of the read were assembled with HiCanu (v2.0, genomeSize=155k) (Nurk et al., 2020). The three resulting HiCanu unitigs were aligned to themselves with Nucmer (v3.1, --maxmatch) (Kurtz et al., 2004) and manually joined to produce a single trimmed and circularized contig. The contig was rotated to start at the same position as the reference. Liftoff was used to annotate the *P. grisea* chloroplast genome (Shumate & Salzberg, 2021).

*P. grisea* mitochondrial contigs were extracted from the polished ONT Flye assembly (see below). To identify mitochondrial contigs, tobacco, pepper, tomato, and eggplant mitochondrial transcript sequences (GenBank IDs NC_006581.1, NC_024624.1, NC_035963.1, and NC_050334.1, respectively) (Sugiyama et al., 2005) were extracted with gffread (G. Pertea & Pertea, 2020) and aligned to the ONT Flye assembly with Minimap2 (v2.17-r941, -x splice). For each query transcriptome, any ONT contig shorter than 500 kbp with at least one alignment at least 100 bp long was considered, and any such contig identified by at least two query transcriptomes was labeled as mitochondrial. These contigs were aligned to the *P. grisea* chloroplast genome which indicated that they were all mitochondrial and not chloroplast sequences. These ONT mitochondrial sequences were aligned to the raw HiCanu contigs (see below) with Nucmer (v3.1, --maxmatch), and nine ONT contigs were manually replaced with two homologous HiCanu contigs. Liftoff was used to annotate the *P. grisea* mitochondrial genome using the *S. melongena* annotation as evidence.

### *P. grisea* genome assembly

*P. grisea* HiFi reads were assembled with HiCanu (v2.0, genomeSize=1500m). *P. grisea* ONT reads at least 38 kbp long and with an average quality score of at least Q12 were assembled with Flye (v2.8.1-b1676, --genome-size 1.5g) (Kolmogorov et al., 2019). The Flye contigs were iteratively polished for two rounds with Freebayes (Garrison & Marth, 2012) . 200,000,000 Illumina short reads (SRA ID SRR7066586) were randomly sampled with seqtk (https://github.com/lh3/seqtk) and aligned to the Flye contigs with BWA-MEM (v0.7.17-r1198-dirty) (H. Li, 2013) . Alignments were sorted and indexed with samtools [(Patro et al., 2017) . Freebayes was used to call variants (v1.3.2-dirty, --skip-coverage 480) and polishing edits were incorporated with bcftools consensus (-i’QUAL>1 && (GT="AA" || GT="Aa")’ -Hla) (Danecek et al., 2021).

The HiCanu contigs were aligned to the *P. grisea* chloroplast and mitochondria genomes with minimap2 (v2.17-r941, -x asm5), and any contigs covered more than 50% by alignments were removed. Potential bacterial contaminant sequence was screened using a process similar to that used by the Vertebrate Genomes Project (Rhie et al., 2021) . The HiCanu contigs were first masked with windowmasker (v1.0.0, -mk_counts -sformat obinary -genome_size 1448242897) (Morgulis et al., 2006). Then, the HiCanu contigs were aligned to all RefSeq bacterial genomes, downloaded on May 21st, 2020, with BLAST (v2.5.0, -task megablast -outfmt "6 std score" - window_masker_db) (Altschul et al., 1990). Any contigs with at least one alignment with an E-value less than 0.0001, a score of at least 100, and a percent-identity of at least 98% were manually inspected, and one contig was removed. To remove potential false haplotypic duplication, HiFi reads were aligned to the screened contigs with Minimap2 (v2.17-r941, -x asm5), and any contigs with at least 50% of the contig with less than 5X coverage were purged (Guan et al., 2020)

The screened and purged contigs were patched with Grafter (https://github.com/mkirsche/Grafter), a beta version of RagTag “patch” (Alonge et al., 2021) . Polished ONT contigs were aligned to the HiCanu contigs with Nucmer (v3.1, -maxmatch -l 100-c 500) and these alignments were used by Grafter to make patches (minq=0 min_weight_supp=10 min_weight=10). Patched contigs were then scaffolded with Bionano optical maps generated at the McDonnell Genome Institute at Washington University. Finally, chromosome-scale scaffolds were manually derived with Hi-C using Juicebox Assembly Tools (Dudchenko et al., 2018). To identify and correct potential misassemblies, HiFi and ONT reads were aligned to the scaffolds with Winnowmap (v1.11, -ax map-pb and -ax map-ont, respectively) and structural variants (SVs) were called with Sniffles (v1.0.12, -d 50 -n -1 -s 3) (Jain, Rhie, Zhang, et al., 2020). We removed any SVs with less than 30% of reads supporting the alternative (ALT) allele and we merged the filtered SV calls with Jasmine (v1.0.10, max_dist=500 spec_reads=3 --output_genotypes) (Kirsche et al., 2021). After merging and manually inspecting the SV calls, a total of four misassemblies were manually corrected. VecScreen did not identify any “strong” or “moderate” hits to the adaptor contamination database (ftp://ftp.ncbi.nlm.nih.gov/pub/kitts/adaptors_for_screening_euks.fa) (https://www.ncbi.nlm.nih.gov/tools/vecscreen/). Finally, we removed any unplaced contigs shorter than 1 kbp. Mercury was used to compute QV and completeness metrics (k=21) (Rhie et al., 2020).

### *P. pruinosa* genome assembly

The *P. pruinosa* genome was assembled just as the *P. grisea* genome, with the following distinctions. HiFi reads were assembled with Hifiasm instead of HiCanu (v0.13-r308, -l0) (Cheng et al., 2021). Also, neither a chloroplast nor mitochondria genome was assembled for *P. pruinosa*. To screen organellar contigs, raw Hifiasm primary contigs were aligned to the *P. pruinosa* reference chloroplast genome (GenBank ID MH019243.1) and the *P. grisea* mitochondria genome. As with *P. grisea*, SVs were called to identify potential misassemblies, and no misassemblies were found in the *P. pruinosa* scaffolds.

### Gene and repeat annotation

Raw RNASeq reads from *P. grisea*, were assessed for quality using FastQC v0.11.9 (*FastQC*, 2015), which were then aligned to the *P. grisea* assembly using STAR aligner (Dobin et al., 2013). Finally, reference-based transcripts were assembled using StringTie v2.1.2 (M. Pertea et al., 2015). We used the portcullis v1.2.0 (Mapleson et al., 2018) method to filter out the invalid splice junctions from the bam alignments. Additionally, we lifted orthologs from the Heinz ITAG4.0 annotation (Hosmani et al., 2019) and the pangenome annotation (L. Gao et al., 2019) using the Liftoff v1.6.1(-exclude_partial -copies) (Shumate & Salzberg, 2021) pipeline. Structural gene annotations were then generated using the Mikado v2.0rc2 (Venturini et al., 2018) framework using the evidence set mentioned above following the Snakemake-based pipeline, [Daijin]. Functional annotation of the Mikado gene models was identified using the blastp alignments to uniprot/swissprot (Bairoch & Apweiler, 2000), TREMBL, Heinz ITAG4.0, and pan genome proteins database (L. Gao et al., 2019; Hosmani et al., 2019) and transferred using the AHRD pipeline (https://github.com/asishallab/AHRD). The *P. pruinosa* assembly was gene-annotated with Liftoff, using the *P. grisea* gene annotation as evidence (-copies). Transposable elements were annotated with EDTA (v1.9.6, --sensitive 1 --anno 1 --evaluate 1 --cds) (Ou et al., 2019) . BUSCO was run on each genome assembly using the “embryophyta_odb10” lineage database (v5.0.0, -e 1e-05 --augustus --long) (Simão et al., 2015).

### Structural variant detection

Structural variation between *P. grisea* and *P. pruinosa* was identified using the same pipeline used to identify structural variant-like mis-assemblies described above. However, instead of aligning *P. grisea* reads to the *P. grisea* assembly and *P. pruinosa* reads to the *P. pruinosa* assembly, P. *grisea* reads were aligned to the *P. pruinosa* assembly and *P. pruinosa* reads were aligned to the *P. grisea* assembly. Also, Winnowmap2 (v2.0) was used instead of Winnowmap for alignments (Jain, Rhie, Hansen, et al., 2020) . SVs intersecting genomic features in **Figure 1G** were counted as previously described (Alonge et al., 2020) based on *P. grisea* annotation v1.3.0

### Tissue collection, RNA extraction, RT-PCR and qRT-PCR

All tissues used were immediately frozen in liquid nitrogen before RNA extraction. For the analysis of *AN1* transcripts in *P. grisea* and *P. pru<u>inosa</u>*, young flower buds were harvested. For *TAGL1* gene expression analysis in the tomato calyx mutant, developing sepals at the open flower stage were harvested. For the analysis of *huskless* (*hu*) and WT sepal gene expression profiles, the first whorl of *hu*, and WT sepals and petals at the stages shown in **Figure 5K** were harvested. Total RNA was extracted with the Zymo Research Quick-RNA Microprep kit following the manufacturer’s protocol. cDNA synthesis was performed using SuperScript IV VILO Master Mix (Thermal Fisher) with 500ng ∼ 1,500ng total RNA input. RT-PCR was performed with KOD One^TM^ PCR Master Mix and primers listed in **Supplemental Table S9**. qRT-PCR was performed using Fast SYBR™ Green Master Mix with primers listed in **Supplemental Table S9** on the Applied Biosystems™ QuantStudio™ 6 system.

### Transcriptome analysis of *huskless* and WT

RNA-seq and differentially expressed genes analyses were performed as previously described with slight modification (Kwon et al., 2022). Briefly, the libraries for RNA-seq were prepared by the KAPA mRNA HyperPrep Kit (Roche). Paired-end 150-base sequencing was conducted on the Illumina sequencing platform (NextSeq, High-Output). Reads for WT and *hu* were trimmed by quality using Trimmomatic (Bolger et al., 2014) (v.0.39, parameters: ILLUMINACLIP:TruSeq3-PE-2.fa:2:40:15:1:FALSE LEADING:30 TRAILING:30 MINLEN:50) and quantified to the reference transcriptome assembly of *P. grisea* v1.3.2 using Salmon v1.4.0 (Patro et al., 2017) . Quantification results from Salmon were imported into R using tximport v1.24.0 (Soneson et al., 2016) . PCA analysis of samples were performed and plotted using DEseq2 v1.36.0 (Love et al., 2014) and pcaExplorer v2.22.0 (Marini & Binder, 2019)with counts of the top 3000 variable genes.

### CRISPR-Cas9 mutagenesis, plant transformation, and selection of mutant alleles

CRISPR-Cas9 mutagenesis was performed following our protocol as previously described (Lemmon et al., 2018; Swartwood & Van Eck, 2019). Briefly, guide RNAs (gRNAs) were designed to be used in the Golden Gate cloning system (all gRNAs used in this study are listed in **Supplemental Table S8**) and were assembled into Level 1 (L1) constructs under the control of the U6 promoter. L1 guide constructs were then assembled with Level 1 constructs pICH47732-NOSpro::NPTII and pICH47742-35S:Cas9 into the binary Level 2 vector pAGM4723. The final binary vectors were then transformed into groundcherry by Agrobacterium tumefaciens-mediated transformation through tissue culture (Swartwood & Van Eck, 2019) . Multiple independent first-generation transgenic plants (T_0_) were genotyped with specific primers surrounding the target sites. T_0_ plants were self-pollinated and the T_1_ generation was genotyped for the target genes and the presence/absence of the CRISPR-Cas9 transgene. We noticed that tissue culture and transformation resulted in a variable frequency of tetraploidy. All mutants were verified as homozygous or biallelic and having only mutant alleles.

### Mapping of the yellow nectar guide variant

The yellow-guide trait displayed classical patterns of Mendelian inheritance of a single recessive gene in the F1 and F2 populations from the cross between *P. grisea* and *P. pruinosa*. A bulk segregant analysis (BSA) was performed using 20 plants from each of the yellow-guide pool and purple-guide pool in the F2 segregating population. All reads were assessed for overall quality by FastQC v0.11.9 (*FastQC*, 2015). Read mapping, variant calling, and SNP-index calculation of the Illumina reads from each pool was done by QTL-seq v2.2.2 (Takagi et al., 2013). Parameters used for the sliding window SNP-index calculation by the qtlplot command were -n1 20 -n2 20 - F 2 -D 250 -d 5 -w 1000 -s 50. The calculated SNP-index in each sliding window was imported into R (R Core Team, 2020) for the final plot.

### EMS mutagenesis and mutant screening in *P. grisea*

A small-scale EMS mutagenesis was performed using ∼1500 (measured by weight) *P. grisea* seeds. Seeds were soaked in distilled water overnight and then treated with 0.2% EMS (ethyl methanesulfonate, Sigma Aldrich) for 6 hours. After treatment, seeds were washed with distilled water thoroughly and sowed into 96-well flats. 4-week-old seedlings were then transplanted into the field. When harvesting, fruits from every four M_1_ plants were bulk harvested into one family. For mutant screening, 80 families of M2s were sowed, transplanted, and screened for sepal related phenotypes.

### Mapping of *huskless*

Three *huskless* phenotype plants were identified from the same family. The pooled DNA from the three mutants, and the pooled DNA from 30 WT-looking siblings from the same family, were obtained by CTAB extraction methods. Libraries were prepared for sequencing using the Kapa Hyper PCR-free Kit and sequenced on Illumina Nextseq (PE150, high output). All reads were assessed for overall quality by FastQC v0.11.9 (*FastQC*, 2015), and trimmed with Trimmomatic v0.39 (Bolger et al., 2014) with parameters ILLUMINACLIP:TruSeq3-PE.fa:2:40:15:1:FALSE LEADING:30 TRAILING:30 MINLEN:75 TOPHRED33. Trimmed paired reads were mapped to the reference *P. grisea* genome using BWA-MEM (H. Li, 2013). Alignments were then sorted with samtools (H. Li et al., 2009) . and duplicates marked with PicardTools (“Picard Toolkit,” 2019). Variants were called with freebayes (Garrison & Marth, 2012) and filtered with VCFtools (Danecek et al., 2011) for SNPs with minimum read depth of 3 and minimum quality value of 20. SNPs that are homozygous in the mutant pool but not homozygous in the WT sibling pool were analyzed for effects on transcripts with snpEff (Cingolani et al., 2012) with *P. grisea* annotation v1.3.0.

### Molecular phylogenetic analyses

In order to determine the phylogenetic relationship between the eleven selected Solanaceae species, eighteen genomes were used to define orthogroups by Conservatory (Hendelman et al., 2021) . Protein sequences of the twenty most conserved orthogroups genes were aligned with MAFFT (v7.487) FFT-NS-2 (Katoh & Standley, 2013) (see **Source Data 5**), before constructing the tree by IQ-tree with the following parameters -st AA -b 100 -pers 0.5 –wbtl (Minh et al., 2020) . For the phylogenetic analysis of AP2-like proteins, protein sequences of the orthologs were retrieved from *P. grisea*, *S. lycopersicum* and *P. axillaris* by BLAST (Altschul et al., 1990). Protein sequences (see **Source Data 6**) were imported in MEGA 11 (Tamura et al., 2021) and aligned with MUSCLE (default parameters). The tree was constructed with Maximum Likelihood method and JTT matrix-based model. Bootstrap values (%) based on 500 replicates are indicated near the branching points; branches below 50% have been collapsed.

### Synteny analysis at the *SlBOB* locus

The scaffold quality of the *P. axillaris* genome in the vicinity of *BOB* is suboptimal so we used SL4.0 with the *P. grisea* genome for the analysis. BLAST search using Petunia *BOB* and *SlBOB* cDNA query sequences against the *P. grisea* genome failed to retrieve a high-confidence hit other than *Phygri09g010120* which is the *BEN* ortholog. BLAST search of genes upstream and downstream of *SlBOB* located their syntenic regions in the *P. grisea* genome. Genomic sequences with annotations from *Solyc10g084240* ∼ *Solyc10g084420*, and from *Phygri10g011780* ∼ *Phygri10g011960* were used in clinker v0.0.23 (Gilchrist & Chooi, 2021) to generate gene translation alignments and visualizations.

## Supplemental data

**The following materials are available in the online version of this article.**

Supplemental Figure 1. Hi-C heatmaps confirm reference assembly structural accuracy.

Supplemental Figure 2. Illustrations of CRISPR-engineered mutations in this study.

Supplemental Figure 3. Maximum Likelihood consensus tree of the euAP2 proteins.

Supplemental Table S1. Genome assembly statistics.

Supplemental Table S2. Annotation stats of *P. gri* and *P. pru* genomes.

Supplemental Table S3. High impact SNP calls of *P. pru* Illumina reads against *P. gri* as reference.

Supplemental Table S4. SVs intersecting CDS.

Supplemental Table S5. SVs intersecting genes.

Supplemental Table 6a: SNPs with predicted high impact on transcripts.

Supplemental Table 6b: Co-segregation test of the G/A SNP in *Phygri09g010120* and the huskless phenotype.

Supplemental Table S7. Genes related to this study.

Supplemental Table S8. CRISPR guides used in this study.

Supplemental File S1. Source Data.

## Acknowledgements

We thank Yuval Eshed and members of the Van Eck, Schatz, and Lippman labs for discussions. We thank R. Santos, B. Semen, and G. Robitaille from the Lippman lab for technical support. We thank A. Horowitz Doyle, K. Swartwood, M. Tjahjadi, L. Randall and P. Keen from the Van Eck laboratory for transformations. We thank T. Mulligan, K. Schlecht, A. Krainer, S. Qiao, and B. Fitzgerald for assistance with plant care. We thank D. Zamir for providing the tomato ICS mimic mutant. We thank Yueqin Yang for assistance with the EMS mutagenesis screen.

## Funding

This work was funded by the Howard Hughes Medical Institute to Z.B.L., and the National Science Foundation Plant Genome Research Program (IOS-1732253) to J.V.E., M.C.S., and Z.B.L.

*Conflict of interest statement*. None declared.

M.C.S., and Z.B.L. conceived, designed, and led the study, and analyzed the data. J.H. led and coordinated the experiments and analyses. M.C.S. and Z.B.L. performed the genome sequencing, M.A. and M.C.S. generated the genome assemblies. S.R. annotated the genomes. M.B. and S.S. prepared DNA for long-read sequencing. S.S. performed the genome sequencing and analysis to identify the tomato ICS mimic mutation. N.T.R. contributed the CRISPR construct targeting PgMPF3. J.V.E. led the CRISPR transformations and generated all the CRISPR T_0_ lines. A.H. contributed to the phylogenetic analyses. J.H., M.A. and Z.B.L. prepared the figures and wrote the manuscript. All authors read, edited and approved the manuscript.

The author responsible for distribution of materials integral to the findings presented in this article in accordance with the policy described in the Instructions for Authors (https://academic.oup.com/plcell) is: Zachary B. Lippman.

## References

Alonge, M., Lebeigle, L., Kirsche, M., Aganezov, S., Wang, X., Lippman, Z. B., Schatz, M. C., & Soyk, S. (2021). Automated assembly scaffolding elevates a new tomato system for high-throughput genome editing. BioRxiv, 2021.11.18.469135. https://doi.org/10.1101/2021.11.18.469135

Alonge, M., Wang, X., Benoit, M., Soyk, S., Pereira, L., Zhang, L., Suresh, H., Ramakrishnan, S., Maumus, F., Ciren, D., Levy, Y., Harel, T. H., Shalev-Schlosser, G., Amsellem, Z., Razifard, H., Caicedo, A. L., Tieman, D. M., Klee, H., Kirsche, M., … Lippman, Z. B. (2020). Major Impacts of Widespread Structural Variation on Gene Expression and Crop Improvement in Tomato. Cell, 182(1), 145–161.e23. https://doi.org/https://doi.org/10.1016/j.cell.2020.05.021

Altschul, S. F., Gish, W., Miller, W., Myers, E. W., & Lipman, D. J. (1990). Basic local alignment search tool. Journal of Molecular Biology, 215(3), 403–410. https://doi.org/https://doi.org/10.1016/S0022-2836(05)80360-2

Añibarro-Ortega, M., Pinela, J., Alexopoulos, A., Petropoulos, S. A., Ferreira, I. C. F. R., & Barros, L. (2022). Chapter Four - The powerful Solanaceae: Food and nutraceutical applications in a sustainable world. In F. Toldrá (Ed.), Advances in Food and Nutrition Research (Vol. 100, pp. 131–172). Academic Press. https://doi.org/https://doi.org/10.1016/bs.afnr.2022.03.004

Bairoch, A., & Apweiler, R. (2000). The SWISS-PROT protein sequence database and its supplement TrEMBL in 2000. Nucleic Acids Research, 28(1), 45–48. https://doi.org/10.1093/nar/28.1.45

Baumann, T. W., & Meier, C. M. (1993). Chemical defence by withanolides during fruit development in Physalis peruviana. Phytochemistry, 33(2), 317–321. https://doi.org/https://doi.org/10.1016/0031-9422(93)85510-X

Bolger, A. M., Lohse, M., & Usadel, B. (2014). Trimmomatic: a flexible trimmer for Illumina sequence data. Bioinformatics, 30(15), 2114–2120. https://doi.org/10.1093/bioinformatics/btu170

Bombarely, A., Moser, M., Amrad, A., Bao, M., Bapaume, L., Barry, C. S., Bliek, M., Boersma, M. R., Borghi, L., Bruggmann, R., Bucher, M., D’Agostino, N., Davies, K., Druege, U., Dudareva, N., Egea-Cortines, M., Delledonne, M., Fernandez-Pozo, N., Franken, P., … Kuhlemeier, C. (2016). Insight into the evolution of the Solanaceae from the parental genomes of Petunia hybrida. Nature Plants, 2(6), 16074. https://doi.org/10.1038/nplants.2016.74

Chang, C., Bowman, J. L., & Meyerowitz, E. M. (2016). Field Guide to Plant Model Systems. Cell, 167(2), 325–339. https://doi.org/https://doi.org/10.1016/j.cell.2016.08.031

Cheng, H., Concepcion, G. T., Feng, X., Zhang, H., & Li, H. (2021). Haplotype-resolved de novo assembly using phased assembly graphs with hifiasm. Nature Methods, 18(2), 170–175. https://doi.org/10.1038/s41592-020-01056-5

Cingolani, P., Platts, A., Wang, L. L., Coon, M., Nguyen, T., Wang, L., Land, S. J., Lu, X., & Ruden, D. M. (2012). A program for annotating and predicting the effects of single nucleotide polymorphisms, SnpEff. Fly, 6(2), 80–92. https://doi.org/10.4161/fly.19695

Danecek, P., Auton, A., Abecasis, G., Albers, C. A., Banks, E., DePristo, M. A., Handsaker, R. E., Lunter, G., Marth, G. T., Sherry, S. T., McVean, G., Durbin, R., & Group, 1000 Genomes Project Analysis. (2011). The variant call format and VCFtools. Bioinformatics, 27(15), 2156–2158. https://doi.org/10.1093/bioinformatics/btr330

Danecek, P., Bonfield, J. K., Liddle, J., Marshall, J., Ohan, V., Pollard, M. O., Whitwham, A., Keane, T., McCarthy, S. A., Davies, R. M., & Li, H. (2021). Twelve years of SAMtools and BCFtools. GigaScience, 10(2), giab008. https://doi.org/10.1093/gigascience/giab008

Deanna, R., Larter, M. D., Barboza, G. E., & Smith, S. D. (2019). Repeated evolution of a morphological novelty: a phylogenetic analysis of the inflated fruiting calyx in the Physalideae tribe (Solanaceae). American Journal of Botany, 106(2), 270–279. https://doi.org/https://doi.org/10.1002/ajb2.1242

Dobin, A., Davis, C. A., Schlesinger, F., Drenkow, J., Zaleski, C., Jha, S., Batut, P., Chaisson, M., & Gingeras, T. R. (2013). STAR: ultrafast universal RNA-seq aligner. Bioinformatics, 29(1), 15–21. https://doi.org/10.1093/bioinformatics/bts635

Dudchenko, O., Shamim, M. S., Batra, S. S., Durand, N. C., Musial, N. T., Mostofa, R., Pham, M., Glenn St Hilaire, B., Yao, W., Stamenova, E., Hoeger, M., Nyquist, S. K., Korchina, V., Pletch, K., Flanagan, J. P., Tomaszewicz, A., McAloose, D., Pérez Estrada, C., Novak, B. J., … Aiden, E. L. (2018). The Juicebox Assembly Tools module facilitates &lt;em&gt;de novo&lt;/em&gt; assembly of mammalian genomes with chromosome-length scaffolds for under $1000. BioRxiv, 254797. https://doi.org/10.1101/254797

*FastQC*. (2015). https://qubeshub.org/resources/fastqc

Gao, H., Li, J., Wang, L., Zhang, J., & He, C. (2020). Transcriptomic variation of the flower–fruit transition in Physalis and Solanum. Planta, 252(2), 28. https://doi.org/10.1007/s00425-020-03434-x

Gao, L., Gonda, I., Sun, H., Ma, Q., Bao, K., Tieman, D. M., Burzynski-Chang, E. A., Fish, T. L., Stromberg, K. A., Sacks, G. L., Thannhauser, T. W., Foolad, M. R., Diez, M. J., Blanca, J., Canizares, J., Xu, Y., van der Knaap, E., Huang, S., Klee, H. J., … Fei, Z. (2019). The tomato pan-genome uncovers new genes and a rare allele regulating fruit flavor. Nature Genetics, 51(6), 1044–1051. https://doi.org/10.1038/s41588-019-0410-2

Garrison, E. P., & Marth, G. T. (2012). Haplotype-based variant detection from short-read sequencing. ArXiv: Genomics.

Gebhardt, C. (2016). The historical role of species from the Solanaceae plant family in genetic research. Theoretical and Applied Genetics, 129(12), 2281–2294. https://doi.org/10.1007/s00122-016-2804-1

Gilchrist, C. L. M., & Chooi, Y.-H. (2021). clinker &amp; clustermap.js: automatic generation of gene cluster comparison figures. Bioinformatics, 37(16), 2473–2475. https://doi.org/10.1093/bioinformatics/btab007

Grandillo, S., & Tanksley, S. D. (1996). QTL analysis of horticultural traits differentiating the cultivated tomato from the closely related species Lycopersicon pimpinellifolium. Theoretical and Applied Genetics, 92(8), 935–951. https://doi.org/10.1007/BF00224033

Guan, D., McCarthy, S. A., Wood, J., Howe, K., Wang, Y., & Durbin, R. (2020). Identifying and removing haplotypic duplication in primary genome assemblies. Bioinformatics, 36(9), 2896–2898. https://doi.org/10.1093/bioinformatics/btaa025

He, C., Münster, T., & Saedler, H. (2004). On the origin of floral morphological novelties. FEBS Letters, 567(1), 147–151. https://doi.org/https://doi.org/10.1016/j.febslet.2004.02.090

He, C., & Saedler, H. (2005). Heterotopic expression of MPF2 is the key to the evolution of the Chinese lantern of Physalis, a morphological novelty in Solanaceae. Proceedings of the National Academy of Sciences of the United States of America, 102(16), 5779–5784. https://doi.org/10.1073/pnas.0501877102

He, C., & Saedler, H. (2007). Hormonal control of the inflated calyx syndrome, a morphological novelty, in Physalis. The Plant Journal, 49(5), 935–946. https://doi.org/https://doi.org/10.1111/j.1365-313X.2006.03008.x

Hendelman, A., Zebell, S., Rodriguez-Leal, D., Dukler, N., Robitaille, G., Wu, X., Kostyun, J., Tal, L., Wang, P., Bartlett, M. E., Eshed, Y., Efroni, I., & Lippman, Z. B. (2021). Conserved pleiotropy of an ancient plant homeobox gene uncovered by cis-regulatory dissection. Cell, 184(7), 1724–1739.e16. https://doi.org/https://doi.org/10.1016/j.cell.2021.02.001

Hosmani, P. S., Flores-Gonzalez, M., van de Geest, H., Maumus, F., Bakker, L. v, Schijlen, E., van Haarst, J., Cordewener, J., Sanchez-Perez, G., Peters, S., Fei, Z., Giovannoni, J. J., Mueller, L. A., & Saha, S. (2019). An improved de novo assembly and annotation of the tomato reference genome using single-molecule sequencing, Hi-C proximity ligation and optical maps. BioRxiv, 767764. https://doi.org/10.1101/767764

Hu, J.-Y., & Saedler, H. (2007). Evolution of the Inflated Calyx Syndrome in Solanaceae. Molecular Biology and Evolution, 24(11), 2443–2453. https://doi.org/10.1093/molbev/msm177

Huang, M., He, J.-X., Hu, H.-X., Zhang, K., Wang, X.-N., Zhao, B.-B., Lou, H.-X., Ren, D.-M., & Shen, T. (2020). Withanolides from the genus Physalis: a review on their phytochemical and pharmacological aspects. Journal of Pharmacy and Pharmacology, 72(5), 649–669. https://doi.org/10.1111/jphp.13209

Itkin, M., Seybold, H., Breitel, D., Rogachev, I., Meir, S., & Aharoni, A. (2009). TOMATO AGAMOUS-LIKE 1 is a component of the fruit ripening regulatory network. The Plant Journal, 60(6), 1081–1095. https://doi.org/https://doi.org/10.1111/j.1365-313X.2009.04064.x

Jain, C., Rhie, A., Hansen, N., Koren, S., & Phillippy, A. M. (2020). A long read mapping method for highly repetitive reference sequences. BioRxiv, 2020.11.01.363887. https://doi.org/10.1101/2020.11.01.363887

Jain, C., Rhie, A., Zhang, H., Chu, C., Walenz, B. P., Koren, S., & Phillippy, A. M. (2020). Weighted minimizer sampling improves long read mapping. Bioinformatics, 36(Supplement_1), i111–i118. https://doi.org/10.1093/bioinformatics/btaa435

Katoh, K., & Standley, D. M. (2013). MAFFT Multiple Sequence Alignment Software Version 7: Improvements in Performance and Usability. Molecular Biology and Evolution, 30(4), 772–780. https://doi.org/10.1093/molbev/mst010

Kim, S., Park, M., Yeom, S.-I., Kim, Y.-M., Lee, J. M., Lee, H.-A., Seo, E., Choi, J., Cheong, K., Kim, K.-T., Jung, K., Lee, G.-W., Oh, S.-K., Bae, C., Kim, S.-B., Lee, H.-Y., Kim, S.-Y., Kim, M.-S., Kang, B.-C., … Choi, D. (2014). Genome sequence of the hot pepper provides insights into the evolution of pungency in Capsicum species. Nature Genetics, 46(3), 270–278. https://doi.org/10.1038/ng.2877

Kirsche, M., Prabhu, G., Sherman, R., Ni, B., Aganezov, S., & Schatz, M. C. (2021). Jasmine: Population-scale structural variant comparison and analysis. BioRxiv, 2021.05.27.445886. https://doi.org/10.1101/2021.05.27.445886

Kolmogorov, M., Yuan, J., Lin, Y., & Pevzner, P. A. (2019). Assembly of long, error-prone reads using repeat graphs. Nature Biotechnology, 37(5), 540–546. https://doi.org/10.1038/s41587-019-0072-8

Kurtz, S., Phillippy, A., Delcher, A. L., Smoot, M., Shumway, M., Antonescu, C., & Salzberg, S. L. (2004). Versatile and open software for comparing large genomes. Genome Biology, 5(2), R12. https://doi.org/10.1186/gb-2004-5-2-r12

Kwon, C.-T., Tang, L., Wang, X., Gentile, I., Hendelman, A., Robitaille, G., Van Eck, J., Xu, C., & Lippman, Z. B. (2022). Dynamic evolution of small signalling peptide compensation in plant stem cell control. Nature Plants, 8(4), 346–355. https://doi.org/10.1038/s41477-022-01118-w

Lemmon, Z. H., Reem, N. T., Dalrymple, J., Soyk, S., Swartwood, K. E., Rodriguez-Leal, D., van Eck, J., & Lippman, Z. B. (2018). Rapid improvement of domestication traits in an orphan crop by genome editing. Nature Plants, 4(10), 766–770. https://doi.org/10.1038/s41477-018-0259-x

Li, H. (2013). Aligning sequence reads, clone sequences and assembly contigs with BWA-MEM. ArXiv: Genomics.

Li, H. (2018). Minimap2: pairwise alignment for nucleotide sequences. Bioinformatics, 34(18), 3094–3100. https://doi.org/10.1093/bioinformatics/bty191

Li, H., Handsaker, B., Wysoker, A., Fennell, T., Ruan, J., Homer, N., Marth, G., Abecasis, G., Durbin, R., & Subgroup, 1000 Genome Project Data Processing. (2009). The Sequence Alignment/Map format and SAMtools. Bioinformatics, 25(16), 2078–2079. https://doi.org/10.1093/bioinformatics/btp352

Li, J., Song, C., & He, C. (2019). Chinese lantern in Physalis is an advantageous morphological novelty and improves plant fitness. Scientific Reports, 9(1), 596. https://doi.org/10.1038/s41598-018-36436-7

Liu, Y., Tikunov, Y., Schouten, R. E., Marcelis, L. F. M., Visser, R. G. F., & Bovy, A. (2018). Anthocyanin Biosynthesis and Degradation Mechanisms in Solanaceous Vegetables: A Review. Frontiers in Chemistry, 6. https://www.frontiersin.org/articles/10.3389/fchem.2018.00052

Love, M. I., Huber, W., & Anders, S. (2014). Moderated estimation of fold change and dispersion for RNA-seq data with DESeq2. Genome Biology, 15(12), 550. https://doi.org/10.1186/s13059-014-0550-8

Lu, J., Luo, M., Wang, L., Li, K., Yu, Y., Yang, W., Gong, P., Gao, H., Li, Q., Zhao, J., Wu, L., Zhang, M., Liu, X., Zhang, X., Zhang, X., Kang, J., Yu, T., Li, Z., Jiao, Y., … He, C. (2021). The Physalis floridana genome provides insights into the biochemical and morphological evolution of Physalis fruits. Horticulture Research, 8(1), 244. https://doi.org/10.1038/s41438-021-00705-w

Mapleson, D., Venturini, L., Kaithakottil, G., & Swarbreck, D. (2018). Efficient and accurate detection of splice junctions from RNA-seq with Portcullis. GigaScience, 7(12), giy131. https://doi.org/10.1093/gigascience/giy131

Marini, F., & Binder, H. (2019). pcaExplorer: an R/Bioconductor package for interacting with RNA-seq principal components. BMC Bioinformatics, 20(1), 331. https://doi.org/10.1186/s12859-019-2879-1

Martínez, M. (1993). The correct application of Physalis pruinosa L. (Solanaceae). TAXON, 42(1), 103–104. https://doi.org/https://doi.org/10.2307/1223312

Meir, Z., Aviezer, I., Chongloi, G. L., Ben-Kiki, O., Bronstein, R., Mukamel, Z., Keren-Shaul, H., Jaitin, D., Tal, L., Shalev-Schlosser, G., Harel, T. H., Tanay, A., & Eshed, Y. (2021). Dissection of floral transition by single-meristem transcriptomes at high temporal resolution. Nature Plants, 7(6), 800–813. https://doi.org/10.1038/s41477-021-00936-8

Minh, B. Q., Schmidt, H. A., Chernomor, O., Schrempf, D., Woodhams, M. D., von Haeseler, A., & Lanfear, R. (2020). IQ-TREE 2: New Models and Efficient Methods for Phylogenetic Inference in the Genomic Era. Molecular Biology and Evolution, 37(5), 1530–1534. https://doi.org/10.1093/molbev/msaa015

Morel, P., Heijmans, K., Rozier, F., Zethof, J., Chamot, S., Bento, S. R., Vialette-Guiraud, A., Chambrier, P., Trehin, C., & Vandenbussche, M. (2017). Divergence of the Floral A-Function between an Asterid and a Rosid Species. The Plant Cell, 29(7), 1605–1621. https://doi.org/10.1105/tpc.17.00098

Morgulis, A., Gertz, E. M., Schäffer, A. A., & Agarwala, R. (2006). WindowMasker: window-based masker for sequenced genomes. Bioinformatics, 22(2), 134–141. https://doi.org/10.1093/bioinformatics/bti774

Muller, G. B., & Wagner, G. P. (1991). Novelty in Evolution: Restructuring the Concept. Annual Review of Ecology and Systematics, 22, 229–256. http://www.jstor.org/stable/2097261

Nurk, S., Walenz, B. P., Rhie, A., Vollger, M. R., Logsdon, G. A., Grothe, R., Miga, K. H., Eichler, E. E., Phillippy, A. M., & Koren, S. (2020). HiCanu: accurate assembly of segmental duplications, satellites, and allelic variants from high-fidelity long reads. Genome Research, 30(9), 1291–1305. https://doi.org/10.1101/gr.263566.120

Ou, S., Su, W., Liao, Y., Chougule, K., Agda, J. R. A., Hellinga, A. J., Lugo, C. S. B., Elliott, T. A., Ware, D., Peterson, T., Jiang, N., Hirsch, C. N., & Hufford, M. B. (2019). Benchmarking transposable element annotation methods for creation of a streamlined, comprehensive pipeline. Genome Biology, 20(1), 275. https://doi.org/10.1186/s13059-019-1905-y

Padmaja, H., Sruthi, S. R., & Vangalapati, M. (2014). INTERNATIONAL JOURNAL OF PHARMACY & LIFE SCIENCES (Int. J. of Pharm. Life Sci.) Review on Hibiscus sabdariffa - A valuable herb.

Pan, I. L., McQuinn, R., Giovannoni, J. J., & Irish, V. F. (2010). Functional diversification of AGAMOUS lineage genes in regulating tomato flower and fruit development. Journal of Experimental Botany, 61(6), 1795–1806. https://doi.org/10.1093/jxb/erq046

Park, S. J., Eshed, Y., & Lippman, Z. B. (2014). Meristem maturation and inflorescence architecture—lessons from the Solanaceae. Current Opinion in Plant Biology, 17, 70–77. https://doi.org/https://doi.org/10.1016/j.pbi.2013.11.006

Paton, A. (1990). A Global Taxonomic Investigation of Scutellaria (Labiatae). Kew Bulletin, 45(3), 399–450. https://doi.org/10.2307/4110512

Patro, R., Duggal, G., Love, M. I., Irizarry, R. A., & Kingsford, C. (2017). Salmon provides fast and bias-aware quantification of transcript expression. Nature Methods, 14(4), 417–419. https://doi.org/10.1038/nmeth.4197

Pertea, G., & Pertea, M. (2020). GFF Utilities: GffRead and GffCompare [version 2; peer review: 3 approved] . F1000Research, 9(304). https://doi.org/10.12688/f1000research.23297.2

Pertea, M., Pertea, G. M., Antonescu, C. M., Chang, T.-C., Mendell, J. T., & Salzberg, S. L. (2015). StringTie enables improved reconstruction of a transcriptome from RNA-seq reads. Nature Biotechnology, 33(3), 290–295. https://doi.org/10.1038/nbt.3122

Picard toolkit. (2019). In Broad Institute, GitHub repository. Broad Institute.

Pnueli, L., Hareven, D., Rounsley, S. D., Yanofsky, M. F., & Lifschitz, E. (1994). Isolation of the Tomato AGAMOUS Gene TAG1 and Analysis of Its Homeotic Role in Transgenic Plants. The Plant Cell, 6(2), 163–173. https://doi.org/10.2307/3869636

Pretz, C., & Deanna, R. (2020). Typifications and nomenclatural notes in Physalis (Solanaceae) from the United States. TAXON, 69(1), 170–192. https://doi.org/https://doi.org/10.1002/tax.12159

R Core Team. (2020). R: A Language and Environment for Statistical Computing. https://www.R-project.org/

Rhie, A., McCarthy, S. A., Fedrigo, O., Damas, J., Formenti, G., Koren, S., Uliano-Silva, M., Chow, W., Fungtammasan, A., Kim, J., Lee, C., Ko, B. J., Chaisson, M., Gedman, G. L., Cantin, L. J., Thibaud-Nissen, F., Haggerty, L., Bista, I., Smith, M., … Jarvis, E. D. (2021). Towards complete and error-free genome assemblies of all vertebrate species. Nature, 592(7856), 737–746. https://doi.org/10.1038/s41586-021-03451-0

Rhie, A., Walenz, B. P., Koren, S., & Phillippy, A. M. (2020). Merqury: reference-free quality, completeness, and phasing assessment for genome assemblies. Genome Biology, 21(1), 245. https://doi.org/10.1186/s13059-020-02134-9

Rydberg, P. A. (1896). The North American species of Physalis and related genera. *New York*: Torrey Botanical Club.

Sato, S., Tabata, S., Hirakawa, H., Asamizu, E., Shirasawa, K., Isobe, S., Kaneko, T., Nakamura, Y., Shibata, D., Aoki, K., Egholm, M., Knight, J., Bogden, R., Li, C., Shuang, Y., Xu, X., Pan, S., Cheng, S., Liu, X., … Fabra, U. P. (2012). The tomato genome sequence provides insights into fleshy fruit evolution. Nature, 485(7400), 635–641. https://doi.org/10.1038/nature11119

Senthil-Kumar, M., & Mysore, K. S. (2011). Caveat of RNAi in Plants: The Off-Target Effect. In H. Kodama & A. Komamine (Eds.), RNAi and Plant Gene Function Analysis: Methods and Protocols (pp. 13–25). Humana Press. https://doi.org/10.1007/978-1-61779-123-9_2

Shenstone, E., Lippman, Z., & van Eck, J. (2020). A review of nutritional properties and health benefits of Physalis species. Plant Foods for Human Nutrition, 75(3), 316–325. https://doi.org/10.1007/s11130-020-00821-3

Shubin, N., Tabin, C., & Carroll, S. (2009). Deep homology and the origins of evolutionary novelty. Nature, 457(7231), 818–823. https://doi.org/10.1038/nature07891

Shumate, A., & Salzberg, S. L. (2021). Liftoff: accurate mapping of gene annotations. Bioinformatics, 37(12), 1639–1643. https://doi.org/10.1093/bioinformatics/btaa1016

Simão, F. A., Waterhouse, R. M., Ioannidis, P., Kriventseva, E. v, & Zdobnov, E. M. (2015). BUSCO: assessing genome assembly and annotation completeness with single-copy orthologs. Bioinformatics, 31(19), 3210–3212. https://doi.org/10.1093/bioinformatics/btv351

Soneson, C., Love, M. I., & Robinson, M. D. (2016). Differential analyses for RNA-seq: transcript-level estimates improve gene-level inferences [version 2; peer review: 2 approved] . F1000Research, 4(1521). https://doi.org/10.12688/f1000research.7563.2

Soyk, S., Lemmon, Z. H., Oved, M., Fisher, J., Liberatore, K. L., Park, S. J., Goren, A., Jiang, K., Ramos, A., van der Knaap, E., van Eck, J., Zamir, D., Eshed, Y., & Lippman, Z. B. (2017). Bypassing Negative Epistasis on Yield in Tomato Imposed by a Domestication Gene. Cell, 169(6), 1142–1155.e12. https://doi.org/10.1016/j.cell.2017.04.032

Spelt, C., Quattrocchio, F., Mol, J., & Koes, R. (2002). ANTHOCYANIN1 of Petunia Controls Pigment Synthesis, Vacuolar pH, and Seed Coat Development by Genetically Distinct Mechanisms. The Plant Cell, 14(9), 2121–2135. https://doi.org/10.1105/tpc.003772

Spelt, C., Quattrocchio, F., Mol, J. N. M., & Koes, R. (2000). anthocyanin1 of Petunia Encodes a Basic Helix-Loop-Helix Protein That Directly Activates Transcription of Structural Anthocyanin Genes. The Plant Cell, 12(9), 1619–1631. https://doi.org/10.1105/tpc.12.9.1619

Sugiyama, Y., Watase, Y., Nagase, M., Makita, N., Yagura, S., Hirai, A., & Sugiura, M. (2005). The complete nucleotide sequence and multipartite organization of the tobacco mitochondrial genome: comparative analysis of mitochondrial genomes in higher plants. Molecular Genetics and Genomics, 272(6), 603–615. https://doi.org/10.1007/s00438-004-1075-8

Swartwood, K., & van Eck, J. (2019). Development of plant regeneration and Agrobacterium tumefaciens-mediated transformation methodology for Physalis pruinosa. *Plant Cell*, Tissue and Organ Culture (PCTOC), 137(3), 465–472. https://doi.org/10.1007/s11240-019-01582-x

Takagi, H., Abe, A., Yoshida, K., Kosugi, S., Natsume, S., Mitsuoka, C., Uemura, A., Utsushi, H., Tamiru, M., Takuno, S., Innan, H., Cano, L. M., Kamoun, S., & Terauchi, R. (2013). QTL-seq: rapid mapping of quantitative trait loci in rice by whole genome resequencing of DNA from two bulked populations. The Plant Journal, 74(1), 174–183. https://doi.org/https://doi.org/10.1111/tpj.12105

Tamura, K., Stecher, G., & Kumar, S. (2021). MEGA11: Molecular Evolutionary Genetics Analysis Version 11. Molecular Biology and Evolution, 38(7), 3022–3027. https://doi.org/10.1093/molbev/msab120

Theißen, G., & Saedler, H. (2001). Floral quartets. Nature, 409(6819), 469–471. https://doi.org/10.1038/35054172

Venturini, L., Caim, S., Kaithakottil, G. G., Mapleson, D. L., & Swarbreck, D. (2018). Leveraging multiple transcriptome assembly methods for improved gene structure annotation. GigaScience, 7(8), giy093. https://doi.org/10.1093/gigascience/giy093

Waterfall, U. T. (1967). PHYSALIS IN MEXICO, CENTRAL AMERICA AND THE WEST INDIES. Rhodora, 69(777), 82–120. http://www.jstor.org/stable/23311644

Waterfall, U. T. (Umaldy T. (1958). A taxonomic study of the genus Physalis in North America north of Mexico. Rhodora, 60, 152–173. https://www.biodiversitylibrary.org/part/124500

Wei, Q., Wang, J., Wang, W., Hu, T., Hu, H., & Bao, C. (2020). A high-quality chromosome-level genome assembly reveals genetics for important traits in eggplant. Horticulture Research, 7(1), 153. https://doi.org/10.1038/s41438-020-00391-0

Weigel, D., & Meyerowitz, E. M. (1994). The ABCs of floral homeotic genes. Cell, 78(2), 203–209. https://doi.org/https://doi.org/10.1016/0092-8674(94)90291-7

Whitson, M. (2012). CALLIPHYSALIS (SOLANACEAE): A NEW GENUS FROM THE SOUTHEASTERN USA. Rhodora, 114(958), 133–147. http://www.jstor.org/stable/23314732

Wilf, P., Carvalho, M. R., Gandolfo, M. A., & Cúneo, N. R. (2017). Eocene lantern fruits from Gondwanan Patagonia and the early origins of Solanaceae. Science, 355(6320), 71–75. https://doi.org/10.1126/science.aag2737

Xu, P., Zhang, Y., Kang, L., Roossinck, M. J., & Mysore, K. S. (2006). Computational Estimation and Experimental Verification of Off-Target Silencing during Posttranscriptional Gene Silencing in Plants. Plant Physiology, 142(2), 429–440. https://doi.org/10.1104/pp.106.083295

Xu, X., Pan, S., Cheng, S., Zhang, B., Mu, D., Ni, P., Zhang, G., Yang, S., Li, R., Wang, J., Orjeda, G., Guzman, F., Torres, M., Lozano, R., Ponce, O., Martinez, D., de la Cruz, G., Chakrabarti, S. K., Patil, V. U., … Centre, W. U. & R. (2011). Genome sequence and analysis of the tuber crop potato. Nature, 475(7355), 189–195. https://doi.org/10.1038/nature10158

Yanofsky, M. F., Ma, H., Bowman, J. L., Drews, G. N., Feldmann, K. A., & Meyerowitz, E. M. (1990). The protein encoded by the Arabidopsis homeotic gene agamous resembles transcription factors. Nature, 346(6279), 35–39. https://doi.org/10.1038/346035a0

Yuste-Lisbona, F. J., Quinet, M., Fernández-Lozano, A., Pineda, B., Moreno, V., Angosto, T., & Lozano, R. (2016). Characterization of vegetative inflorescence (mc-vin) mutant provides new insight into the role of MACROCALYX in regulating inflorescence development of tomato. Scientific Reports, 6(1), 18796. https://doi.org/10.1038/srep18796

Zamora-Tavares, M. del P., Martínez, M., Magallón, S., Guzmán-Dávalos, L., & Vargas-Ponce, O. (2016). Physalis and physaloids: A recent and complex evolutionary history. Molecular Phylogenetics and Evolution, 100, 41–50. https://doi.org/https://doi.org/10.1016/j.ympev.2016.03.032

Zhang, J.-S., Li, Z., Zhao, J., Zhang, S., Quan, H., Zhao, M., & He, C. (2014). Deciphering the Physalis floridana Double-Layered-Lantern1 Mutant Provides Insights into Functional Divergence of the GLOBOSA Duplicates within the Solanaceae . Plant Physiology, 164(2), 748–764. https://doi.org/10.1104/pp.113.233072

Zhang, W.-N., & Tong, W.-Y. (2016). Chemical Constituents and Biological Activities of Plants from the Genus Physalis. Chemistry & Biodiversity, 13(1), 48–65. https://doi.org/https://doi.org/10.1002/cbdv.201400435

Zhao, J., Tian, Y., Zhang, J.-S., Zhao, M., Gong, P., Riss, S., Saedler, R., & He, C. (2013). The euAP1 protein MPF3 represses MPF2 to specify floral calyx identity and displays crucial roles in Chinese lantern development in Physalis. The Plant Cell, 25(6), 2002–2021. https://doi.org/10.1105/tpc.113.111757

